# KIM-1-mediated anti-inflammatory activity is preserved by MUC1 induction in the proximal tubule during ischemia-reperfusion injury

**DOI:** 10.1101/2021.05.20.445053

**Authors:** Mohammad M. Al-bataineh, Carol L. Kinlough, Zaichuan Mi, Edwin K. Jackson, Stephanie M. Mutchler, David R. Emlet, John A. Kellum, Rebecca P. Hughey

**Affiliations:** Renal-Electrolyte Division, Department of Medicine, University of Pittsburgh School of Medicine, Pittsburgh, PA; Department of Pharmacology and Chemical Biology, University of Pittsburgh School of Medicine, Pittsburgh, PA; Center for Critical Care Nephrology, Department of Critical Care Medicine. University of Pittsburgh School of Medicine, Pittsburgh, PA

**Author notes:** **CORRESPONDENCE**: Mohammad Al-bataineh, DVM, MS, PhD, Assistant Professor of Medicine, Renal-Electrolyte Division, School of Medicine, University of Pittsburgh, Scaife Hall S931, 3550 Terrace Street, Pittsburgh, PA 15261. **Supplemental Material available at DOI**: https://doi.org/10.6084/m9.figshare.14364890.

## Abstract

Cell-associated kidney injury molecule-1 (KIM-1) exerts an anti-inflammatory role following kidney injury by mediating efferocytosis and downregulating the NF-κB pathway. KIM-1 cleavage blunts its anti-inflammatory activities. We reported that Mucin 1 (MUC1) is protective in a mouse model of ischemia-reperfusion injury (IRI). As both KIM-1 and MUC1 are induced in the proximal tubule (PT) during IRI and are ADAM17 substrates, we tested the hypothesis that MUC1 protects KIM-1 activity. *Muc1* KO mice and wild-type (WT) littermates were subjected to IRI. KIM-1, MUC1 and ADAM17 levels (and signaling pathways) were assessed by immunoblotting. PT localization was assessed by confocal microscopy and *in situ* proximity ligation assay. Findings were extended using human kidneys and urine, and KIM-1-mediated efferocytosis assays in mouse PT cultures. In response to tubular injury in mouse and human kidneys, we observed induction and co-expression of KIM-1 and MUC1 in the PT. Compared to WT, *Muc1* KO mice had higher urinary KIM-1 and lower kidney KIM-1. KIM-1 was apical in PT of WT kidneys, but predominately with luminal debris in *Muc1* KO mice. Efferocytosis was reduced in *Muc1* KO PT cultures when compared to WT cells, while inflammation was increased in *Muc1* KO kidneys when compared to WT mice. MUC1 was cleaved by ADAM17 in PT cultures, and blocked KIM-1 shedding in MDCK cells. We conclude that KIM-1-mediated efferocytosis and thus anti-inflammatory activity during IRI is preserved in the injured kidney by MUC1 inhibition of KIM-1 shedding.

**NEW & NOTEWORTHY:** KIM-1 plays a key role in the recovery of the tubule epithelium during renal IRI by mediating efferocytosis and associated signaling that suppresses inflammation. Excessive cleavage of KIM-1 by ADAM17 provides decoy receptor that aggravates efferocytosis and subsequent signaling. Our data from studies in mice, patients and cultured cells show that MUC1 is also induced during IRI and competes with KIM-1 for cleavage by ADAM17. Consequently, MUC1 protects KIM-1 anti-inflammatory activity in the damaged kidney.

## INTRODUCTION

Kidney injury molecule-1 (KIM-1) is a type I transmembrane glycoprotein that is rapidly induced at the gene and protein levels after acute kidney injury (AKI) (1, 2). Specifically, KIM-1 is upregulated on the apical surface of surviving proximal tubule (PT) epithelial cells of the kidney following ischemic and toxic acute kidney injury (AKI) (1-3). KIM-1 is also upregulated in chronic kidney disease, renal cell carcinoma, and polycystic kidney disease (4-6). Mechanistically, it was recently reported that the transcription factor signal transducer and activator of transcription 3 (STAT3) is phosphorylated by checkpoint kinase 1 (ChK1) and/or extracellular signal-regulated kinases 1 and 2 (ERK1/2) following kidney injury which leads to upregulation of KIM-1 expression (7, 8). Structurally, KIM-1 contains an extracellular Ig-like domain and mucin-like domain, one transmembrane domain and a cytoplasmic domain with a tyrosine kinase phosphorylation motif (9). The ectodomain of human and mouse KIM-1 is cleaved by a disintegrin and metalloproteinase domain 17 (ADAM17) generating soluble KIM-1 (cleaved form), and thereby constitutively shed into the urine providing an early and sensitive biomarker for kidney injury (10, 11). There are also reports that cleavage by ADAM10 may contribute to shedding of mouse and human KIM-1 (12).

Cell-associated KIM-1 exhibits an anti-inflammatory function and thus plays a protective role after kidney injury by (i) downregulating the proinflammatory NF-κB signaling pathway in a phosphoinositide 3-kinase (PI3K)-dependent manner, and by (ii) acting as a receptor for phosphatidylserine (an “eat me” signal presented by apoptotic cells) (1, 2). The cell-associated form of KIM-1 enables PT epithelial cells to recognize and mediate phagocytosis of apoptotic and necrotic cells (efferocytosis) following kidney injury, which in turn prevents inflammation and promotes kidney repair (1, 2). However, accelerated shedding of KIM-1 that is mediated by ADAM17 and regulated by p38 mitogen-activated protein kinase (p38 MAPK) in a cell culture model, competitively inhibits efferocytosis, as the excess soluble KIM-1 acts as a decoy to the cell-associated KIM-1 (10, 13).

The involvement of ADAM17 in a variety of processes reflects the wide range of ADAM17 substrates including the pro-inflammatory cytokine TNF-α, epidermal growth factor receptor (EGFR) ligand, KIM-1, and MUC1 (10, 14). Despite the substantial clinical impact of KIM-1 cleavage by ADAM17 during AKI, there is no *in vivo* data about the regulation of KIM-1 shedding following AKI. It is known that ADAM17 is weakly expressed by normal kidney, while its expression is significantly upregulated in PT, capillaries, and glomeruli after AKI (15, 16). But, with more than 80 different reported ADAM17 substrates, it is clear that targeting or blocking cleavage of KIM-1 would involve a cell-specific event.

There is emerging evidence that the transmembrane glycoprotein Mucin 1 (MUC1), expressed on the apical surface of polarized epithelial cells in kidney, plays novel and important roles under physiological and pathological conditions (17). We previously reported that MUC1 plays a protective role in a mouse model of ischemia-reperfusion injury (IRI) by stabilizing HIF-1α and β-catenin levels and downstream signaling (18, 19). Although MUC1 is normally expressed in the thick ascending limb and distal nephron, we discovered that MUC1 is present at low levels in the PT and significantly induced after ischemic injury (17, 19). There are also reports that the MUC1 ectodomain is shed in response to a specific cleavage by ADAM17 but not ADAM10 (14, 20).

As both KIM-1and MUC1 are protective during IRI, while both are induced in the PT, we were intrigued in preliminary studies to find increased KIM-1 during IRI in the urine of *Muc1* KO mice when compared to WT littermates. We initially expected to find increased KIM-1 expression in response to increased injury in the absence of MUC1, but instead we found reduced KIM-1 levels in the kidney. As activation of upstream pathways regulating KIM-1 expression was no different in *Muc1* KO and WT mice, we also assessed p38 MAPK regulation of ADAM17 expression but found no differences. However, when we compared data from mouse kidneys and urine during IRI, and cell culture models of KIM-1 and MUC1 shedding, we established that MUC1 stabilizes KIM-1 and its activity in the PT by direct competition as a substrate for cleavage by ADAM17. This study provides new insights into our understanding of the mechanism of KIM-1 shedding *in vivo* that has a significant clinical impact on kidney recovery following injury. It also provides a novel aspect of MUC1 protection of the kidney during IRI by preserving KIM-1 function.

## MATERIALS AND METHODS

### Cells and Reagents

Madin-Darby Canine Kidney (MDCK 2001) cells were obtained from Kai Simons in the European Molecular Biology Laboratory (EMBL) in Heidelberg, Germany. These cells were grown in DMEM/F12 (D6421) supplemented with 5% FBS (GIBCO/ThermoFisher, Grand Island, NY) and maintained at 37°C in 5% CO_2_. All chemical compounds used in the studies presented here were purchased from Sigma-Aldrich (St. Louis, MO, USA) unless otherwise stated. The sources and other information of primary antibodies are detailed in supplementary Table 1 (DOI: https://doi.org/10.6084/m9.figshare.14364890). FITC- or Rhodamine-labeled antibodies and TO-PRO Cy5 were obtained from Molecular Probes (ThermoFisher, Grand Island, NY). Secondary horseradish peroxidase (HRP)-conjugated anti-rabbit immunoglobulin G (IgG) and anti-mouse IgG antibodies were obtained from SeraCare (Milford, MA, USA). HRP-conjugated anti-β-actin antibodies were obtained from Proteintech (Rosemont, IL, USA).

### Preparation of Primary Cultures of Proximal Tubule Cells (PTCs)

Primary cultures of mouse kidney PTCs from *Muc1* KO, WT littermates, and transgenic mice expressing human MUC1 were generated as previously described (1, 21, 22). Briefly, the kidney cortex was dissected and minced into pieces (∼1 mm) in prewarmed Hank’s Balanced Salt Solution (HBSS; Life Technologies, Grand Island, NY). Tissues were then gently washed to remove debris and digested in HBSS containing 1 mg/ml collagenase type II (Sigma-Aldrich, St. Louis, MO). Tissue dissociation was performed for 60 min in a shaker (200 rpm) at 37°C with intermittent mixing (vortex for 30 sec). The collagenase reaction was terminated using 5% BSA, and cell preparations were then washed and passed through a cell strainer (nylon mesh) with 40-μm pores to remove undigested material and glomeruli. Finally, cells were washed twice and resuspended in collagen I-coated glass coverslips or 12-well plates in a medium composed of equal volumes of DMEM and Ham’s F-12 plus 60 nM sodium selenate, 5 mg/ml transferrin, 2 mM glutamine, 50 nM dexamethasone, 1 nM triiodothyronine, 10 ng/ml epidermal growth factor, 5 mg/ml insulin, 20 mM D-glucose, 2% (vol/vol) FBS, 20 mM HEPES, and antibiotics, pH 7.4 (reagents from Life Technologies and Sigma-Aldrich). The epithelial cells were used in experiments between 6 and 7 days of culture without passaging. All solute concentrations are wt/vol unless noted otherwise.

### Animals and IRI Protocol

All studies were conducted in accordance with the National Institutes of Health Guide for the Care and Use of Laboratory Animals and approved by the University of Pittsburgh School of Medicine Animal Care and Use Committee. All experiments were conducted using 12-16-week-old male and female C57BL/6J mice. *Muc1* global KO mice on a C57BL/6J background were as previously described and have no obvious phenotype under normal physiological conditions other than increased sensitivity to bacterial infections (23, 24). Acute kidney injury (AKI) was induced as previously described using the two-kidney hanging-weight model of renal ischemia-reperfusion injury (IRI) (25). Briefly, each mouse was anesthetized with isoflurane (5% for induction and 1%–2% for maintenance) in oxygen and placed in the supine position on a thermostatically controlled heating system connected to a rectal temperature sensor to maintain body temperature at 37°C. Ischemia was induced by passing a 4/0 silk suture between the renal artery and vein of a kidney and the ends of the suture were passed over a pulley system. Next, the ends were attached to 1-gram weights and the weights are allowed to hang, thus compressing the renal artery with the suture. After 20 minutes of ischemia, the suture was removed, and the procedure was repeated on the opposite kidney followed by reperfusion for 48 hr. Sham mice were treated exactly like injured mice except that the weights were not attached to the ends of the suture.

After 48 h recovery, mean arterial blood pressure (MABP) and heart rate (HR) were measured after inserting a PE-10 catheter into the carotid artery of anesthetized mouse and connected to a digital blood pressure analyzer (Micro-Med, Inc., Louisville, KY). Renal blood flow (RBF) was measured by placing a transit-time flow probe (0.5SB with J-reflector and no handle; custom-designed by Transonic Systems, Inc.; Ithaca, NY) on either the right or left renal artery of the anesthetized mouse and connected to a model T402-PP Transonic flowmeter.

Glomerular filtration rate (GFR) was estimated as described previously (Ref. #25). Briefly, a constant rate infusion (10 μl/min) of ^14^C-inulin (0.0035 μCi/min dissolved in 0.9% saline containing 2.45% albumin) was initiated through the femoral vein catheter. While for collection of urine, a short segment of PE-50 tubing was inserted into silastic tubing, a small hole was made in the rostral end of the bladder using a cautery, and the silastic-covered PE-50 tubing was inserted into the bladder. After completion of surgery, a one-hour rest period was allowed to permit the mouse to stabilize and to let plasma ^14^C-inulin levels approach steady state. At the end of the one-hour rest period, urine was collected for one hour, and the mouse is carefully moved to the prone position. ^14^C-inulin was determined in urine and plasma by measuring radioactivity in these samples, and GFR was estimated by calculating the renal clearance of ^14^C-inulin. KIM-1 was also measured in urine as a marker for PT injury using a mouse ELISA kit (Abcam, Cambridge, MA; ab119596). As a marker of distal nephron injury, neutrophil gelatinase-associated lipocalin (NGAL) was also measured in urine with a mouse ELISA kit (Abcam; ab119601). At the end of the experiment, a blood sample was collected in heparin from the carotid artery catheter, and Serum creatinine (sCr) was measured at the University of Texas Southwestern Medical Center using a Capillary Electrophoresis method (26).

### Kidney Tissue Preparation and Confocal Immunofluorescence Microscopy

One kidney from each mouse was sliced lengthwise and placed in 4% DEPC-treated paraformaldehyde, methacarn (chloroform/methanol/acetic acid) for histology or was snap frozen in liquid nitrogen and then stored at -80°C for subsequent preparation of homogenates for further analysis (e.g. immunoblotting). Subsets of kidneys in 4% paraformaldehyde were sent to the University of Pittsburgh’s pathology laboratory (the P30 O’Brien Core) for preparation of paraffin blocks and for production of kidney slides. Immunofluorescence labeling was subsequently performed, after removing paraffin and antigen retrieval, using antibodies against mouse KIM-1 raised in goat (1:200 dilution), followed by a secondary donkey anti-goat antibody coupled to Alexa 488 (1:400 dilution). Tissues were also co-labeled with an antibody raised in rabbit against the organic anion transporter 1 (OAT1) localized on the PT basolateral surface (1:400 dilution) followed by a secondary antibody raised in donkey coupled to CY3 (1:800 dilution). Mouse kidney tissues were also labeled with a monoclonal antibody raised in Armenian hamster against MUC1 cytoplasmic tail (CT2, 1:200 dilution), followed by a secondary goat anti-Armenian hamster antibody coupled to Alexa 488 (1:400 dilution). Immunolabeled tissues were mounted in *VECTASHIELD*^®^ mounting medium (Vector Laboratories) and imaged in a confocal laser scanning microscope (Leica TCS SP5, Model upright DM 6000S, Leica Microsystems Inc., Buffalo Grove, IL, USA) using a 63x objective with identical laser settings for all samples.

### Immunoblotting

These experiments were performed according to published methods (19, 27) and adapted as follows. The snap-frozen kidney quarter (∼0.4 mg) was homogenized in ice-cold lysis buffer (All solute concentrations are wt/vol unless noted otherwise): 150 mM NaCl, 10 mM Tris (pH 7.4), 1% Triton X-100, 0.5% Igepal, 1 mM EDTA, 1 mM ethylene glycol-bis(β-aminoethyl ether)-N,N,N′,N′-tetraacetic acid (EGTA), and both protease and phosphatase inhibitor cocktails (EMD Millipore, Burlington, MA, USA). The protein samples were then separated by SDS-PAGE on a 4–12% gradient gel (Nu-PAGE, Invitrogen), followed by transfer to nitrocellulose membranes (0.45 μm, Millipore). Then, the blots were probed with anti-MUC1 CT2 antibody (1:1000 dilution), antibodies raised in rabbit against: ADAM17 (1:1000 dilution), p-STAT3 (1:1000 dilution), ERK1/2 (1:5000 dilution), p-ERK1/2 (1:5000 dilution); p38 MAPK (1:5000 dilution), p-p38 MAPK (1:5000 dilution), NFκB p65 (1:5000 dilution), and TLR-4 (1:5000 dilution), or antibodies raised in goat against the mouse TIM-1/KIM-1/HAVCR (1:1000 dilution), or mouse antibody against the STAT3 (1:1000 dilution), or anti-β-actin antibody raised in mouse (1:20,000 dilution) to confirm equal loading of protein. Each probed protein was then visualized and quantified using Bio-Rad Clarity ECL, a Bio-Rad ChemiDoc Imager and Bio-Rad Quantity One or Image Lab software. Blots of samples from *Muc1* KO and WT mouse kidneys were developed side-by-side for the same time period.

Isolation of nuclear and cytoplasmic fractions of soluble proteins from a quarter of a kidney was performed using the NE-PER Nuclear and Cytoplasmic Extraction Kit as per manufacturer’s instructions (ThermoFisher Scientific, Waltham, MA, USA). Aliquots of the nuclear and cytoplasmic fractions were subjected to immunoblotting for NFκB p65 (1:5000 dilution), and β-actin (cytoplasmic fraction) or Histone H3 (nuclear fraction). Blots of samples from *Muc1* KO and wild-type mouse kidneys were developed side-by-side for the same time.

### Quantitative, real-time, reverse-transcriptase polymerase chain reaction

Whole kidney mRNA was isolated using Trizol. One microgram of total RNA was added to a cDNA synthesis kit (BioRad) and cDNA was added to reactions containing SYBR green master mix and a pairing of the following primers: KIM1<u>[RT1]</u> forward 5’-TTC AGG AAG CTG AGC AAA CAT-3’, KIM1 reverse 5’-CCCCAACATGTCGTTGTGATT-3’, 18s forward 5’-GTAACCCGTTGAACCCCATT-3’, 18S reverse 5’-CCATCCAATCGGTAGTAGCG-3’. Differences in expression were calculated using the 2^-DDCt^ method.

### Human Kidneys with Ischemic Damage and Urine Samples

Whole adult human kidneys were obtained from the Center for Organ Recovery and Education (CORE, Pittsburgh, PA) through protocol #462 approved by the University of Pittsburgh Committee for Oversight of Research and Clinical Training Involving Decedents. Kidney samples were from donation-after-cardiac-death or brain-dead donors that were not accepted for transplant. Therefore, we anticipated that these kidneys exhibit some degree of ischemic damage, allowing the opportunity to study the expression of KIM-1, the most highly upregulated protein in the PT after kidney injury. Supplementary Table 3 (DOI: https://doi.org/10.6084/m9.figshare.14364890) lists the wedge biopsy reports of the samples used in this study. All reports list some degree of inflammation, tubular injury, or fibrosis (albeit mild) and further supports our anticipation that these kidneys underwent some sort of stress conditions that would induce KIM-1 expression. Kidneys were then processed as previously described (21). For human kidney, tissues were processed and 5 µm-thick cryosections were prepared as described above. Immunofluorescence labeling was performed using the mouse monoclonal antibody B27.29 against human MUC1 tandem repeat (1:1000 dilution) with goat antibodies against human KIM-1 (1:200 dilution), followed by a secondary donkey anti-mouse antibody coupled to CY3 (1:800 dilution), and donkey anti-goat antibody coupled to Alexa 488 (1:400 dilution), respectively. The B27.29 epitope maps to two separate parts of the tandem repeats including the PDTRP peptide and the glycosylated Thr* and Ser* (GVT*S*APDTRPAPG) and therefore affinity can be affected by glycosylation. As human MUC1 has 40-100 tandem repeats this antibody is much more sensitive for IF staining than the anti-cytoplasmic tail antibody CT2. Immunolabeled tissues were mounted and imaged as described above.

### *In Situ* Proximity Ligation Assay

Proximity ligation assay (PLA) was performed according to the manufacturer’s instructions (Duolink In Situ detection reagent-Red, Sigma Aldrich). Briefly, kidney tissues were permeabilized with 0.5% Triton X-100 in PBS (pH 7.4) and incubated overnight with the mouse monoclonal antibody B27.29 against MUC1 extracellular domain tandem repeat (1:1000 dilution), and with rabbit antibodies against ADAM17 (1:100 dilution). Next day, donkey anti-mouse PLA^minus^, and donkey anti-rabbit PLA^plus^ probes were applied, followed by ligation and amplification with Duolink detection reagent Texas Red according to the manufacturer’s instructions (Duolink In Situ detection reagent-Red, Sigma Aldrich). The samples were then imaged in a confocal laser scanning microscope (Leica TCS SP5, Model upright DM 6000S, Leica Microsystems Inc., Buffalo Grove, IL, USA) using a 20x objective with identical laser settings for all samples. All PLA assays were performed in two independent experiments with a negative control where only ADAM17 antibody was added.

### *In Vitro* Cell Culture Model of KIM-1 Shedding

MDCK cells were transfected with pcDNA 3.1(-) neo mKIM-1 encoding a cDNA for mouse KIM-1, and a stable mixed population of MDCK-KIM-1 cells was obtained by growth in G418. MDCK-KIM-1 cells plated in a 12-well dish were transiently transfected the next day with human *MUC1* cDNA (pcDNA3.1 *MUC1*) or empty vector (EV). The next day, cells were incubated for 2 h in serum free media with either 1.6 μM phorbol-12-myristate-13-acetate in DMSO (PMA; induces ADAM17 metalloprotease-mediated shedding of KIM-1 and MUC1), 10 mM BB-94 in DMSO (Batimastat (Abcam, Cambridge, MA), a broad-spectrum matrix metalloprotease inhibitor), both PMA and BB-94, or vehicle (DMSO). Media was collected on ice and then centrifuged for 5 min at 2000 x g. The supernatant was moved to a clean tube and incubated overnight with 0.5 μg of either goat anti-mouse KIM-1 or mouse anti-MUC1 (B27.29) antibodies and Protein G conjugated to Sepharose 4B (ThermoFisher). Beads were washed with 0.5 ml HEPES-buffered saline (HBS, 10 mM HEPES-HCl, pH 7.4, 150 mM NaCl) with 1% Triton X-100 and then 0.5 ml HBS before heating in Bio-Rad sample buffer with β-mercaptoethanol. Cells were washed twice with ice-cold Dulbecco’s phosphate buffered saline (Corning Cellgro, Manassas, VA) before extraction with 0.2 ml detergent solution (100 mM NaCal, 40 mM KCl, 1 mM EDTA, 20 mM HEPES-KCl, pH 7.4, 10% glycerol, 1% NP40, 0.4% deoxycholate) containing protease inhibitors (EMD Millipore, Burlington, MA, USA). Cell extract (2.5%) or immunoprecipitates of KIM-1 or MUC1 from media were immunoblotted for KIM-1 or MUC1 using SDS-PAGE on a Bio-Rad TGX 4-15% acrylamide gradient gel and transfer to nitrocellulose (0.45 μm, Millipore). Blots were developed with Bio-Rad Clarity Max Western ECL Substrate and imaged with a Bio-Rad Chemidoc and Bio-Rad Image Lab software.

### *In Vitro* Primary Cell Culture Model of MUC1 Shedding

Primary cultures of proximal tubule cells from transgenic mice expressing human MUC1 (n=3) were grown for one week on plastic, then treated with either 3.0 μM PMA or vehicle (DMSO) in serum free media for 1 h in the presence or absence of protease inhibitors against both ADAM10 and ADAM17 (GW280264; 1.0 μM) or specific for ADAM10 (GI254023; 0.5 μM) (28). Inhibitor concentrations were based on the published IC50 values for GW280264 and GI254023 with ADAM17 and ADAM10 (28). Others have reported mouse KIM-1 shedding from HEK293 cells based on inhibition by GI254023 (4 μM) (29) although the IC50 for ADAM10 is 5 nM (28). Cell extract or media were immunoblotted for MUC1 as described above. Transgenic mice expressing human MUC1 were used because no antibodies exist to study mouse extracellular domain shedding. The only available antibody for mouse MUC1 is the Armenian hamster CT2 antibody that recognizes the MUC1 cytoplasmic tail.

### Preparation of Apoptotic Cells

Apoptotic thymocytes were prepared as previously described (1, 10, 30). Briefly, primary mouse thymocytes were disrupted into a single-cell suspension by passing through a 40-μm cell strainer. Next day, thymocytes were labeled with carboxyfluorescein succinimidyl ester (CFSE; CellTrace CFSE dye, Invitrogen, Eugene, OR, USA) according to manufacturer’s instructions and exposed to UV radiation (254 nm) for one minute to induce apoptosis before incubation overnight at 37°C in 5% CO2. Using annexin V/Dead Cell Apoptosis Kit (Alexa Fluor 488 annexin V/Dead Cell Apoptosis Kit with Alexa Fluor 488 annexin V and propidium iodide, Invitrogen, Eugene, OR, USA), we observed more than 90% of exposed cells were deemed to be early apoptotic cells as evidence of Annexin V staining (binds to phosphatidylserine; PS: an “eat me” signal), whereas less than 5% of those were positive for the uptake of propidium iodide.

### Phagocytosis Assay

To quantitate phagocytosis of apoptotic cells, we used a previously established method (1, 10, 30). Briefly, primary cultures of mouse kidney PTCs obtained from *Muc1* KO and WT littermates were grown on glass coverslips or 12-well plate and then incubated with 2 × 10^6^ CFSE fluorescently labeled (FITC) apoptotic thymocytes for one hour at 37°C. After washing vigorously five times with ice-cold PBS (without Mg^+2^/Ca^+2^) to remove non-ingested apoptotic cells, PTC monolayers on glass coverslips were fixed, premetallized, and co-stained as described above. To quantify phagocytic uptake of apoptotic cells (efferocytosis), total number of internalized apoptotic cells per total number of epithelial cells were scored by blinded microscopic counting of random fields (five images per mouse; n = 4 mice). PTC monolayers grown on 12-well plate were solubilized using a lysis buffer containing 62.5 mM EDTA pH 8.0, 50 mM Tris pH 8.0, 4 mg sodium deoxycholate, and 1% NP-40 (IGEPAL). Fluorescence of the lysates was then analyzed using a multi detection plate reader (Promega Glomax Multi Detection System, Madison, WI). PTC monolayers not incubated with thymocytes were considered as a reference control.

### Statistical Analyses

All analyses were performed using GraphPad Prism version 9.0.0 (GraphPad Software, San Diego, California USA). Comparisons involving only two groups were conducted with unpaired *t*-test, while multiple groups were analyzed using two-way ANOVA with a Tukey post hoc test. *P* < 0.05 was considered significant and presented with an asterisk(s). All data are reported as mean values ±SE.

## RESULTS

### MUC1 Protection of Kidney Function During IRI is Associated with a Role in the Proximal Tubule

*Muc1* KO mice and wild-type (WT) littermates were subject to 20 min ischemia and 48 h recovery using the kidney hanging-weight protocol, or sham surgery. While heart rate, mean arterial blood pressure and renal blood flow were not different in *Muc1* KO mice and WT littermates after IRI or sham surgery (Fig. S1A-C; DOI: https://doi.org/10.6084/m9.figshare.14364890), we did find a significant reduction in the glomerular filtration rate (GFR) in the *Muc1* KO mice after IRI when compared to *Muc1* KO sham mice (Fig. 1A). We also found a significant increase in serum creatinine in the *Muc1* KO mice after IRI when compared to *Muc1* KO sham mice or WT mice after IRI or sham surgery (Fig. 1B). Moreover, we found that the proximal tubule (PT) marker of kidney injury, KIM-1, but not the distal tubule injury marker of kidney injury, NGAL, was increased in the urine of *Muc1* KO mice after IRI when compared to *Muc1* KO mice sham and WT mice after IRI (Fig. 1C-D). Altogether, these data indicate that MUC1 protection during IRI is not related to changes in kidney hemodynamics, but reflects a role in the renal epithelial cells of the proximal tubules where MUC1 induction was previously observed (17, 18).

**Figure 1.**
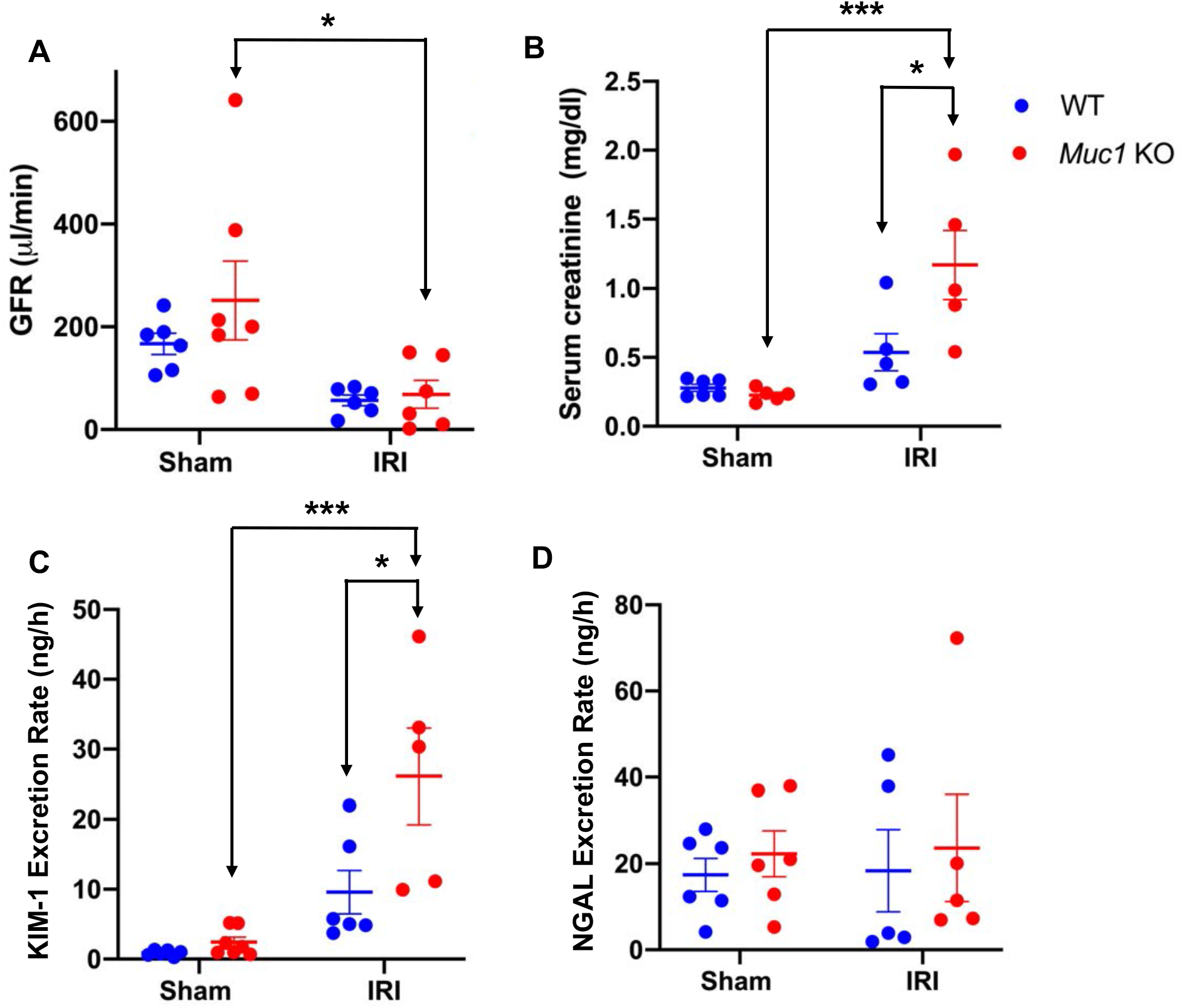
MUC1 protection of kidney function during IRI is associated with a role in the proximal tubule. Kidneys of *Muc1* KO mice (red) and wild-type (WT) littermates (blue) were subjected to 20 min ischemia and 48 h recovery using the kidney hanging-weight protocol (IRI) or sham surgery (Sham). Prior to sacrifice, mice were assessed for (A) glomerular filtration rate (GFR). Blood and urine were collected at sacrifice and assayed for (B) serum creatinine, (C) urinary KIM-1 excretion rate, and (D) urinary NGAL excretion rate. Mean ± SE are shown. Significant differences by two-way ANOVA are noted (*: p<0.05; ***: p<0.001, n=5-7).

Using our established confocal immunofluorescence (IF) microscopy protocols (27, 31), we observed a strong induction and co-localization of KIM-1 and MUC1 in the same PT of WT mouse kidneys subjected to IRI (Fig. 2A). Similar results were observed by co-staining sections from human kidneys with ischemic damage (Fig. 2C & S2). Indeed, we observed induction of both KIM-1 and MUC1 in the same PT mainly at the apical membrane with substantial co-localization.

**Figure 2.**
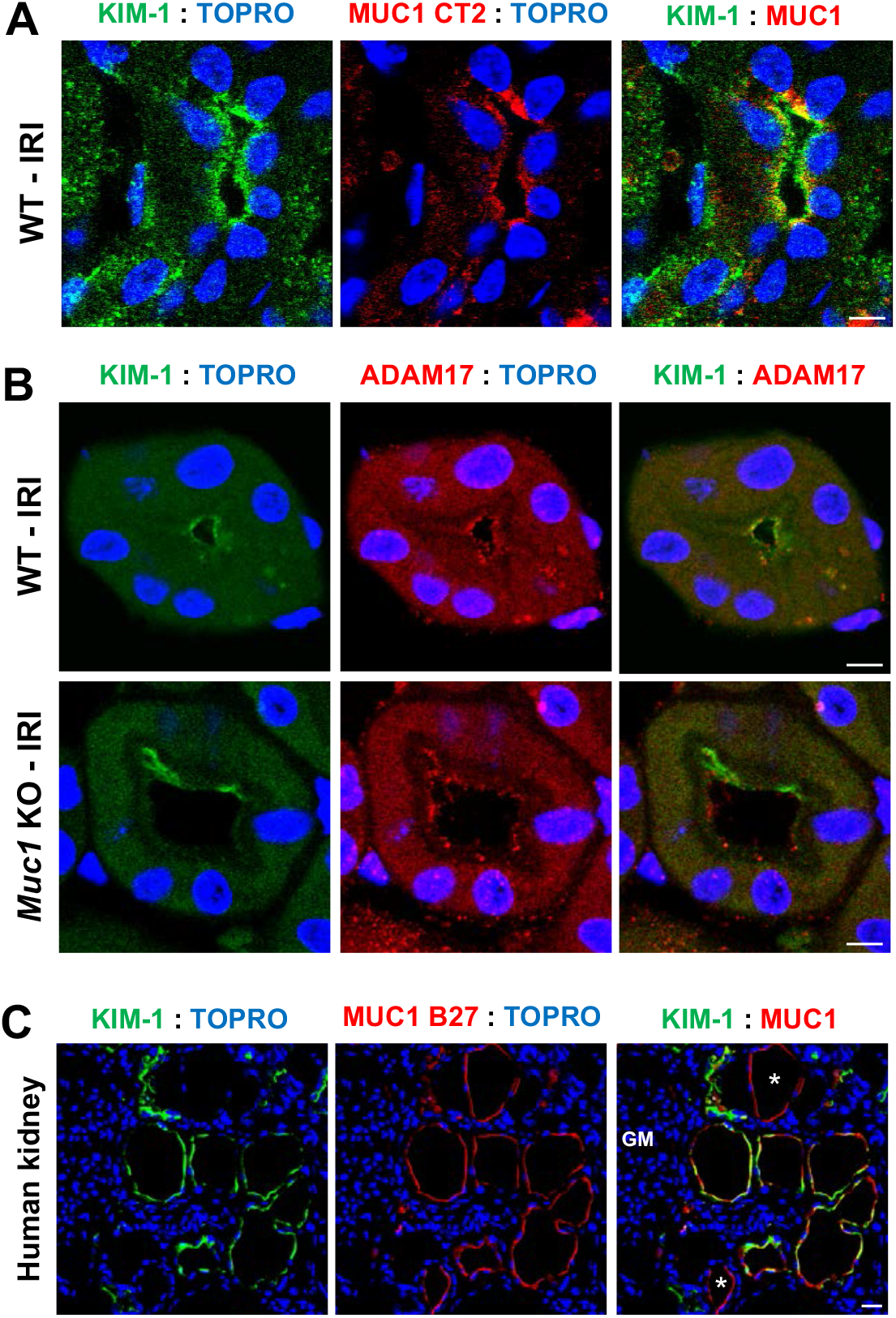
KIM-1 and MUC1 are induced and co-expressed in the proximal tubule in the setting of ischemia. Kidneys of *Muc1* KO mice and wild-type (WT) littermates were subjected to 20 min ischemia and 48 h recovery (IRI) or sham surgery (Sham). Kidneys were harvested at sacrifice and stained for (A) KIM-1 (green), and MUC1 (CT2, red) and nuclei (TOPRO, blue), or (B) KIM-1 (green), and ADAM17 (red) and nuclei (TOPRO, blue), as indicated. (C) Human kidneys with ischemic damage were stained for KIM-1 (green), MUC1 (B27.29, red) and nuclei (TOPRO, blue) as indicated. MUC1 staining was also observed in tubules not expressing KIM-1, consistent with MUC1 expression in TAL, DCT and CD. Scale bar = 10 μm (n=3 mice).

### Kidney KIM-1 Levels are Significantly Lower in *Muc1* KO Mice than WT Mice after IRI

The increased urinary KIM-1 observed in the absence of MUC1 during IRI could reflect an increase in KIM-1 expression in the PT in response to more severe injury. Surprisingly, immunoblotting kidney extracts after IRI revealed a significant two-fold lower level of KIM-1 in *Muc1* KO mice when compared to WT mice (Fig. 3A) while MUC1 levels increased 40% during IRI (Fig. 3B). Interestingly, we found a significant upregulation of the levels of kidney KIM-1 mRNA in both *Muc1* KO and WT littermates following IRI when compared to sham surgery, yet we did not observe a significant difference based on genotype (Fig. 3C). We considered several mechanisms for this observation: (i) MUC1 could sequester ADAM17 from KIM-1 after injury but we did not observe a difference in ADAM17 subcellular localization in *Muc1* KO mouse kidney compared to WT littermate kidneys after injury, as we noticed co-localization between KIM-1 and ADAM17 in both WT and *Muc1* KO mouse kidneys (Fig. 2B). (ii) MUC1 could stimulate the ERK1/2 → STAT3 → KIM-1 pathway reported to increase KIM-1 during IRI, but we found no changes in activation/phosphorylation of ERK1/2 and STAT3 by immunoblotting suggesting posttranslational effects of MUC1 on KIM-1 expression (Fig. S3A-B; DOI: https://doi.org/10.6084/m9.figshare.14364890). (iii) Zhang et al. previously reported that ADAM17-mediated cleavage of KIM-1 is accelerated by p38 MAP kinase activation (11), but we found no difference in the levels of active phosphorylated p38 MAPK nor ADAM17 by immunoblotting kidney extracts from WT or *Muc1 KO* mice during IRI (Fig. S4A-B; DOI: https://doi.org/10.6084/m9.figshare.14364890). Our results indicate that the pathway of KIM-1 accelerated shedding was apparently activated in our mouse model of IRI, but this induction of ADAM17 activity was not affected by MUC1 levels.

**Figure 3.**
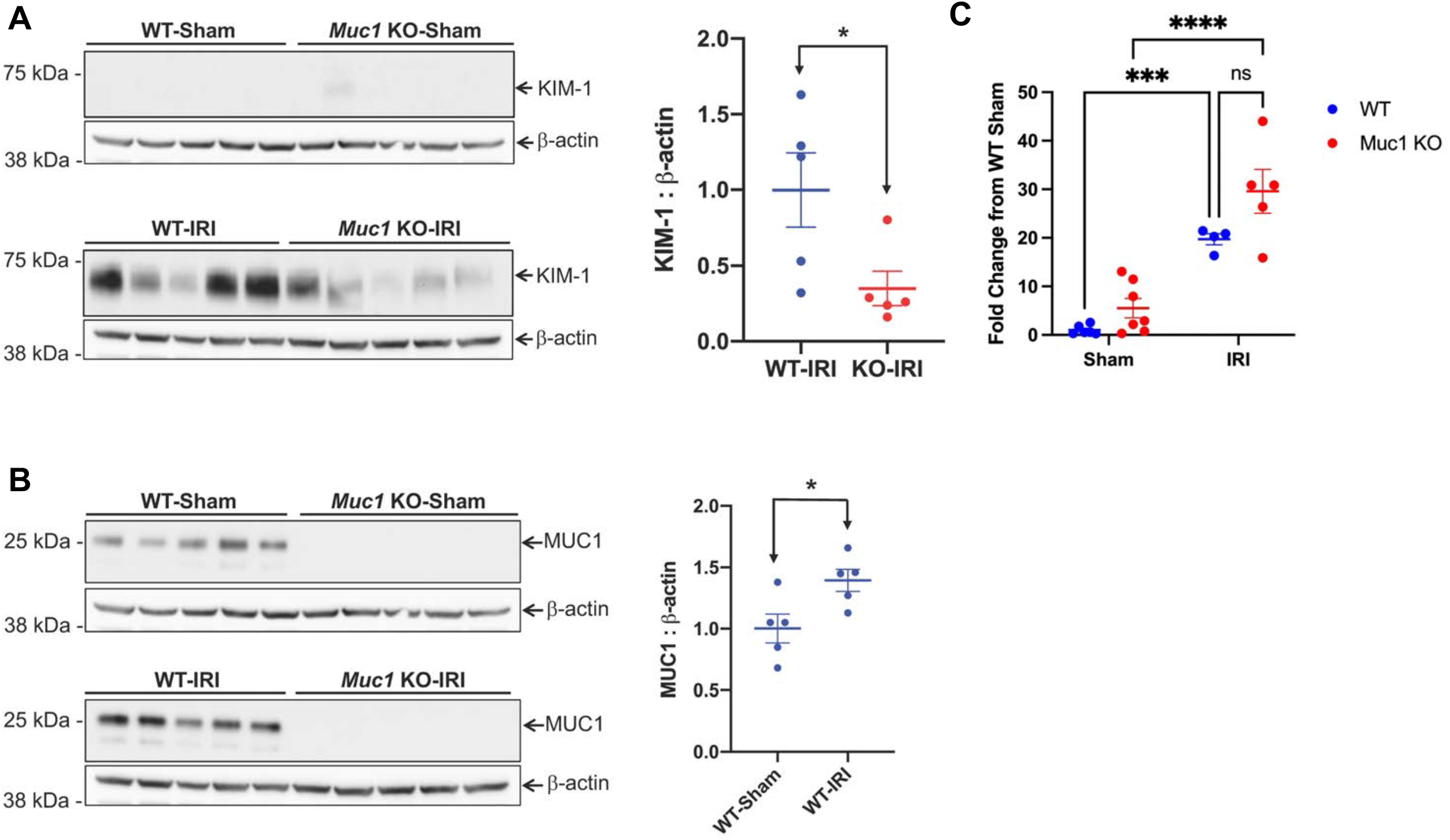
Kidney KIM-1 expression is significantly lower in *Muc1* KO mice than WT mice after IRI. Kidneys of *Muc1* KO mice (red) and wild-type (WT) littermates (blue) were subjected to 20 min ischemia and 48 h recovery (IRI) or sham surgery (Sham). Kidneys were harvested at sacrifice and extracts of homogenates were subjected to immunoblotting for (A) KIM-1 and β-actin or (B) MUC1 small subunit (using CT2) and β-actin. Levels of KIM-1 or MUC1 were normalized to β-actin, and values for WT-IRI mice were set at 1. Data are presented as mean ± SE. Significant differences by Student *t-test* are noted (*; p<0.05, n=5). (C) Levels of KIM-1 mRNA were assessed by RT-qPCR and relative levels presented with WT sham set at 1. Data were analyzed by two-way ANOVA and significant differences are noted (***: p<0.001; ****: p<0.0001, n=5-7).

### MUC1, ADAM17, and KIM-1 are in the Same Complex in Kidney in the Setting of Ischemia

Thathiah et al. previously demonstrated that MUC1 and ADAM17 are in the same protein complex in human uterine epithelial cells(14). To assess this interaction in kidney, we used both confocal IF microscopy and an *in situ* proximity ligation assay (PLA) (27) (32). First, we observed an induction and IF co-localization of both MUC1 and ADAM17 within the same PT of kidneys from WT mice subjected to IRI (Fig. 4A), and human kidneys with ischemic damage (Fig. 4B). Although we could not detect a signal for MUC1 in the glomeruli, we observed a strong staining for ADAM17 in the glomeruli (Fig. 4B, middle) which we used as one of our negative controls for the PLA. We also observed a robust PLA signal for MUC1 and ADAM17 in human kidney with ischemic injury that was completely absent in the negative control lacking the anti-MUC1 antibody and absent in the glomerulus that lacks MUC1 expression (Fig. 4C). As there is no PLA probe available for the Armenian hamster CT2 antibody that recognizes the mouse MUC1 we were not able to assess this interaction in mouse kidneys. Importantly, we also observed co-localization between KIM-1, MUC1 and ADAM17 within the same PT of human kidneys with ischemic damage (Fig. 4D). Altogether, these data further suggest that KIM-1, MUC1 and ADAM17 are in the same protein complex, and ADAM17 likely mediates KIM-1 and MUC1 shedding in kidney.

**Figure 4.**
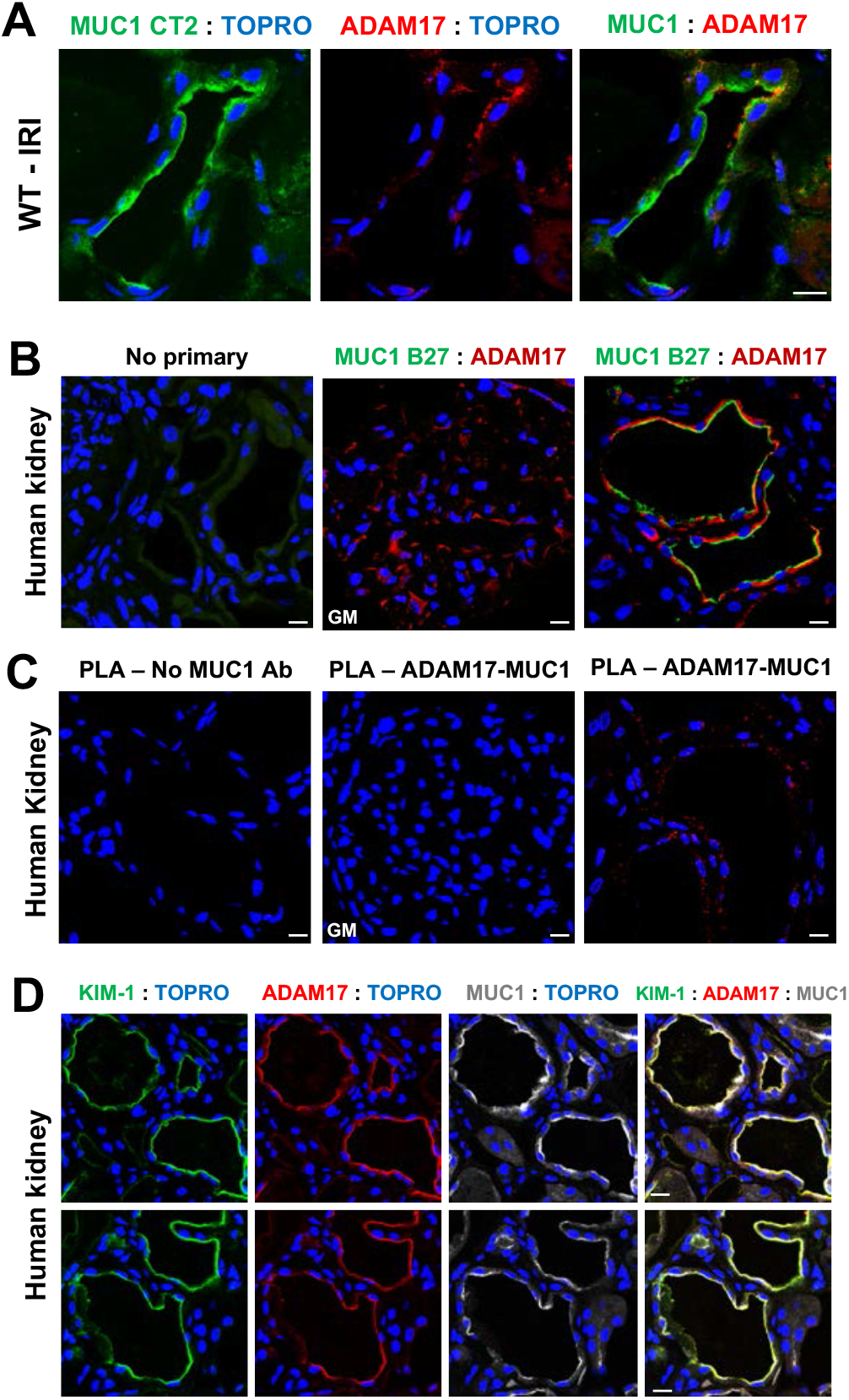
MUC1, ADAM17, and KIM-1 are in the same complex in the kidney in the setting of ischemia. (A) Kidneys of wild-type (WT) mice were subjected to 20 min ischemia and 48 h recovery. Kidneys were harvested at sacrifice and stained for MUC1 small subunit (CT2, green), ADAM17 (red) and nuclei (TOPRO, blue) as indicated. (B) Human kidneys with ischemic damage incubated with no primary antibodies (left) or with antibodies against MUC1 extracellular tandem repeats (B27.29, green), ADAM17 (red) and nuclei (TOPRO, blue) as indicated. (C) *In situ* PLA (proximity ligation assay) was carried out using proximity probes against MUC1 (mouse B27.29) and ADAM17 (rabbit pAb) to visualize MUC1/ADAM17 complex in human kidney sections. Notice that the PLA signal is absent in the negative control lacking the anti-MUC1 antibody, and absent in the glomerulus (GM) that lacks MUC1 expression despite high levels of ADAM17 expression. (D) Human kidneys with ischemic damage were co-stained for KIM-1 (green), ADAM17 (red), MUC1 extracellular tandem repeats (B27.29, gray), and nuclei (TOPRO, blue) as indicated. Scale bar = 10 μm (n=3 mice).

### KIM-1 Shedding is Reduced by MUC1 Expression in a Cell Culture Model

Using primary cultures of PT cells prepared from transgenic mice expressing human MUC1, we did observe ADAM-17-dependent shedding of MUC1 as previously described in human uterine epithelial cells(14) (Fig. S5; DOI: https://doi.org/10.6084/m9.figshare.14364890). To test the hypothesis that MUC1 competes with KIM-1 as a substrate for ADAM17, we used an *in vitro* model of cultured Madin-Darby Canine Kidney (MDCK) cells stably transfected with mouse KIM-1 and transiently transfected with human *MUC1* cDNA or empty vector. The following day, cells were incubated in serum-free medium with either (i) PMA (an ADAM17 activator), (ii) BB-94 (a broad spectrum matrix metalloprotease inhibitor), (iii) both PMA and BB-94, or (iv) vehicle (DMSO). By immunoblotting cell extracts and immunoprecipitates from culture medium, we observed that KIM-1 shedding was significantly enhanced after treating cells with PMA and inhibited after BB-94 treatment (Fig. 5). Levels of KIM-1 shedding (∼1% per h) were similar to a previous study in cultured cells (10). Using two-way ANOVA, we found that KIM-1 shedding was significantly reduced overall by MUC1 expression (p<0.01) (Fig. 5C). We conclude that MUC1 and KIM-1 compete as substrates for ADAM17.

**Figure 5.**
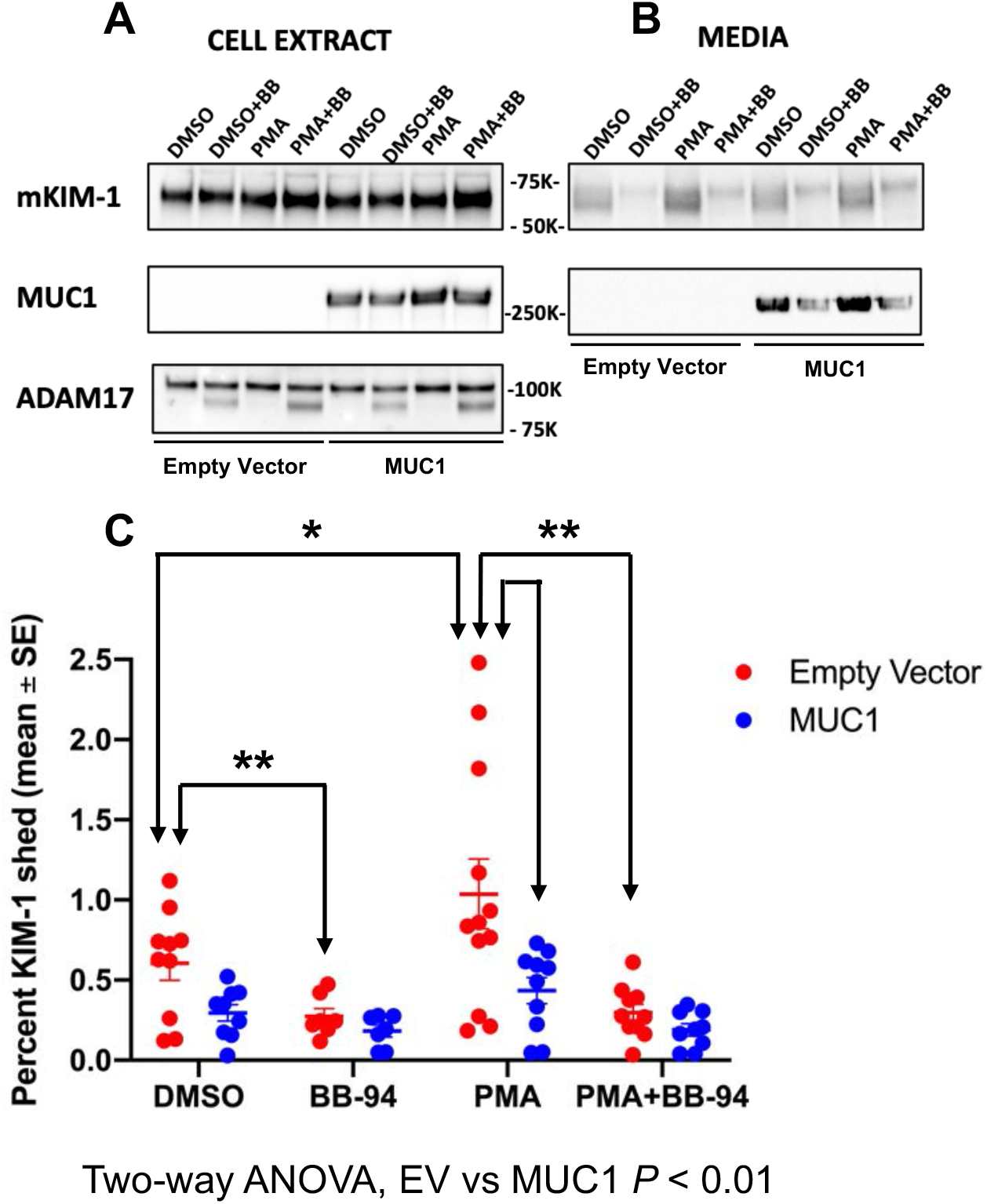
KIM-1 shedding is reduced by MUC1 expression in a cell culture model. MDCK cells stably transfected with mouse KIM-1 were transiently transfected with human MUC1 cDNA or Empty Vector (EV). The next day, cells were incubated for 2 h in serum free media with either (i) 1.6 mM phorbol-12-myristate-13-acetate (PMA) to induce ADAM17 metalloprotease-mediated shedding of KIM-1 and MUC1, (ii) 10 mM BB-94 (a broad-spectrum matrix metalloprotease inhibitor), (iii) both PMA and BB-94, or (iv) vehicle (DMSO). (A) Cell extract (2.5%) or (B) immunoprecipitates of KIM-1 or MUC1 from media were immunoblotted for KIM-1 or MUC1 large subunit (B27.29). (C) Levels of KIM-1 in the medium are presented as percent of levels in the same cells transfected with MUC1 (blue) or empty vector (red). Data were analyzed by two-way ANOVA and significant differences are noted (*: p<0.05; **: p<0.01, n=7-12).

### Cell Association and Function of KIM-1 are Reduced after IRI in the Absence of MUC1

Using confocal IF, we observed KIM-1 staining in the WT mouse kidney PT localized primarily to the apical membrane (Fig. 6A), with a low level of KIM-1 staining surrounding cellular debris within the adjacent tubule lumen consistent with efficient efferocytosis (Fig. 6A; insert). In contrast, KIM-1 staining is more diffuse in *Muc1* KO mouse kidney PT following injury (Fig. 6B), and KIM-1 staining is primarily associated with cell debris in the lumen of the PT (Fig. 6B; insert). The data are consistent with enhanced shedding of KIM-1 from the PT after IRI in the absence of MUC1.

**Figure 6.**
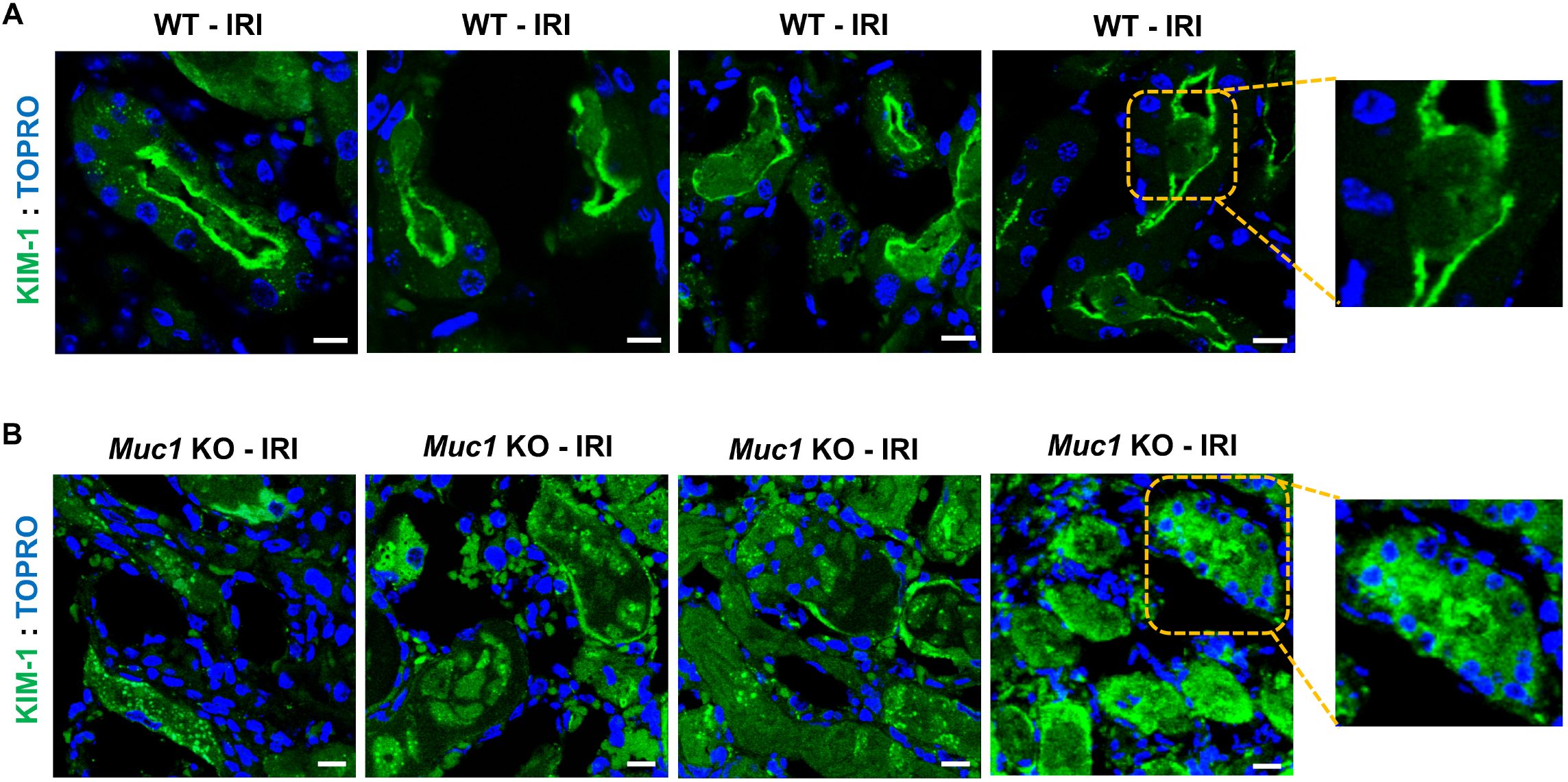
Cell-associated KIM-1 is reduced after IRI in the absence of MUC1. Kidneys of *Muc1* KO mice and wild-type (WT) littermates were subjected to 20 min ischemia and 48 h recovery (IRI). Kidneys (n=4) were harvested at sacrifice and stained for KIM-1 (green) and nuclei (TOPRO, blue). Note that KIM-1 in WT kidneys (A) is primarily cell associated or surrounds luminal cell debris (see insert), while KIM-1 staining in *Muc1* KO kidneys (B) is more diffuse and found primarily with cell debris (see insert) (n=4).

As KIM-1-mediated efferocytosis is dependent on cell-associated KIM-1, we used previously established assays (1, 2, 30) to measure uptake of fluorescently labeled (FITC) apoptotic thymocytes by primary cultures of PT cells (Fig. 7). We observed a significant 22% reduction in thymocyte uptake by *Muc1* KO PTCs compared to WT PTCs by IF (Fig. 7A-B) (p < 0.003). Using a plate reader, *Muc1* KO PTCs growing on plastic exhibited significantly less fluorescence (76%) (Fig. 7C); when compared to WT PTCs (p < 0.01). And, immunoblotting cell extracts revealed lower levels of KIM-1 expression in *Muc1* KO (71%) when compared to WT PTCs (Fig. 7D-E). Thymocytes were also observed in KIM-1-expressing cells but not in KIM-1-negative cells (Fig. 7F and 7G). Taken together, the decreased efferocytosis in *Muc1* KO PTCs could be explained, in part, by reduced surface expression of KIM-1 in the absence of MUC1.

**Figure 7.**
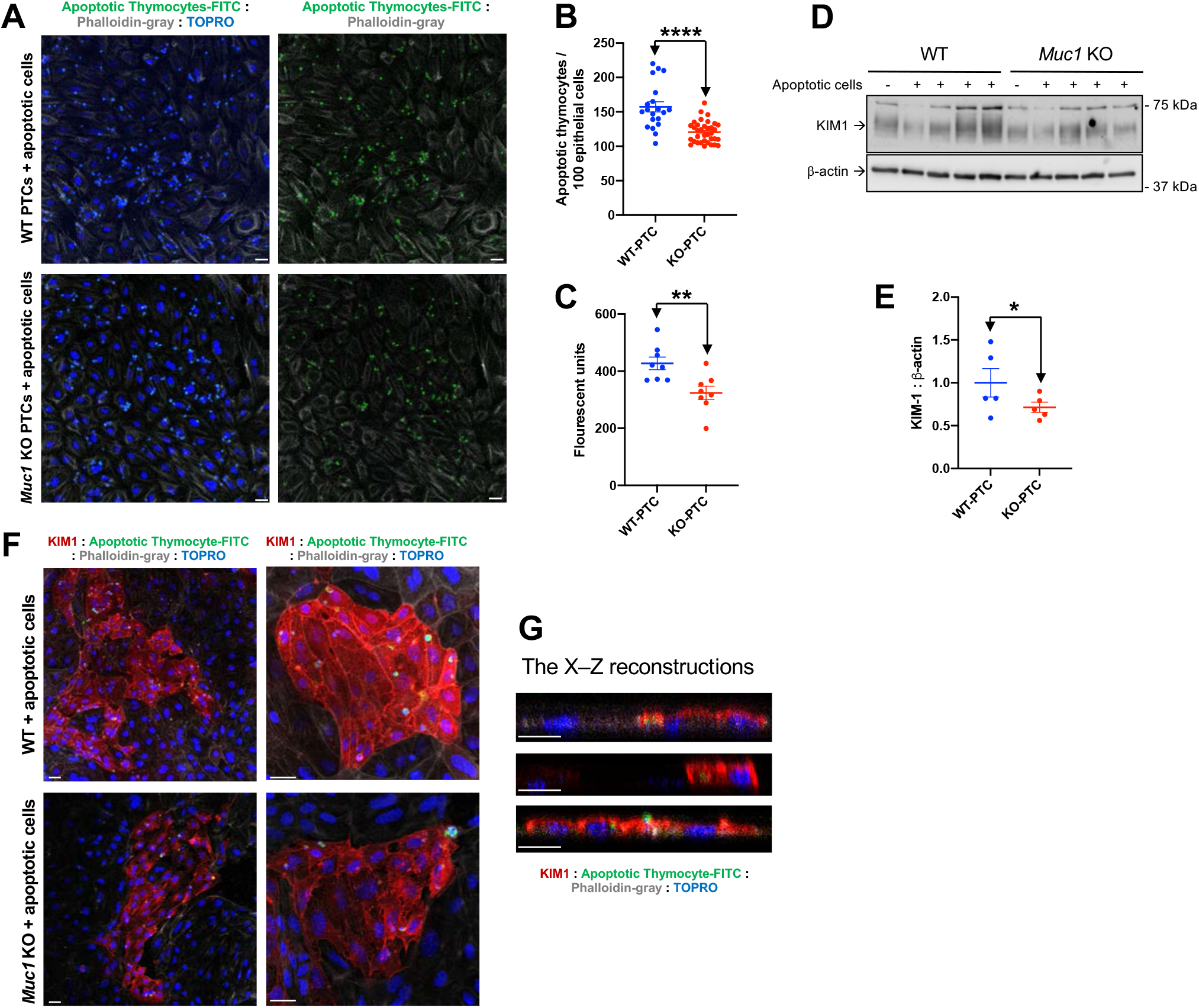
Decreased efferocytosis in primary cultured proximal tubule cells from *Muc1* KO compared to WT littermates. Primary cultures of mouse kidney proximal tubule cells (PTCs) obtained from *Muc1* KO (red) and WT littermates (blue) were grown on glass coverslips (A, B, F and G) or 12-well plates (C-E) and then incubated with CFSE fluorescently-labeled (green) apoptotic thymocytes. (A) Co-cultured PT monolayers with FITC-thymocytes (green) on coverslips were co-stained for actin (phalloidin, gray) and nuclei (TOPRO, blue). Scale bar = 10 μm. (B) Total number of internalized apoptotic cells per total number of epithelial cells were scored by blinded microscopic counting of random fields. (C) Co-cultured PTC monolayers with FITC-thymocytes on 12-well plates were washed, solubilized and fluorescence of the lysates was quantitatively assessed by plate reader. (D) PTC lysates were subjected to immunoblotting for KIM-1 and β-actin, levels of KIM-1 were normalized to β-actin, and values for WT mice were set at 1. (E) Data from (D) are presented as mean ± SE. Significant differences by Student *t-test* are noted (*; p<0.05, n=4). (F) Co-cultured PC monolayers with FITC-thymocytes (green) were co-stained for KIM-1 (red), phalloidin (gray), and nuclei (TOPRO, blue). (G) Confocal reconstruction (x-z) of KIM-1 and FITC-thymocytes demonstrates KIM-1 localization at the phagocytic cup and surrounding phagocytosed apoptotic cells within PTCs. Scale bar = 10 μm (n=4).

Cell-associated KIM-1 exhibits an anti-inflammatory function by downregulating Toll-Like Receptor 4 (TLR4) and its downstream signaling proinflammatory NF-κB pathway following injury (2). By immunoblotting kidney extracts we observed a significantly higher increase in TLR4, and NF-κB (nuclear and cytoplasmic), during IRI in *Muc1 KO* mice than WT mice (Fig. 8A-C). Altogether, the data indicate that MUC1 stabilizes cell-associated KIM-1 during IRI which could protect the kidney by downregulating innate immunity and inflammation (Fig. 9).

**Figure 8.**
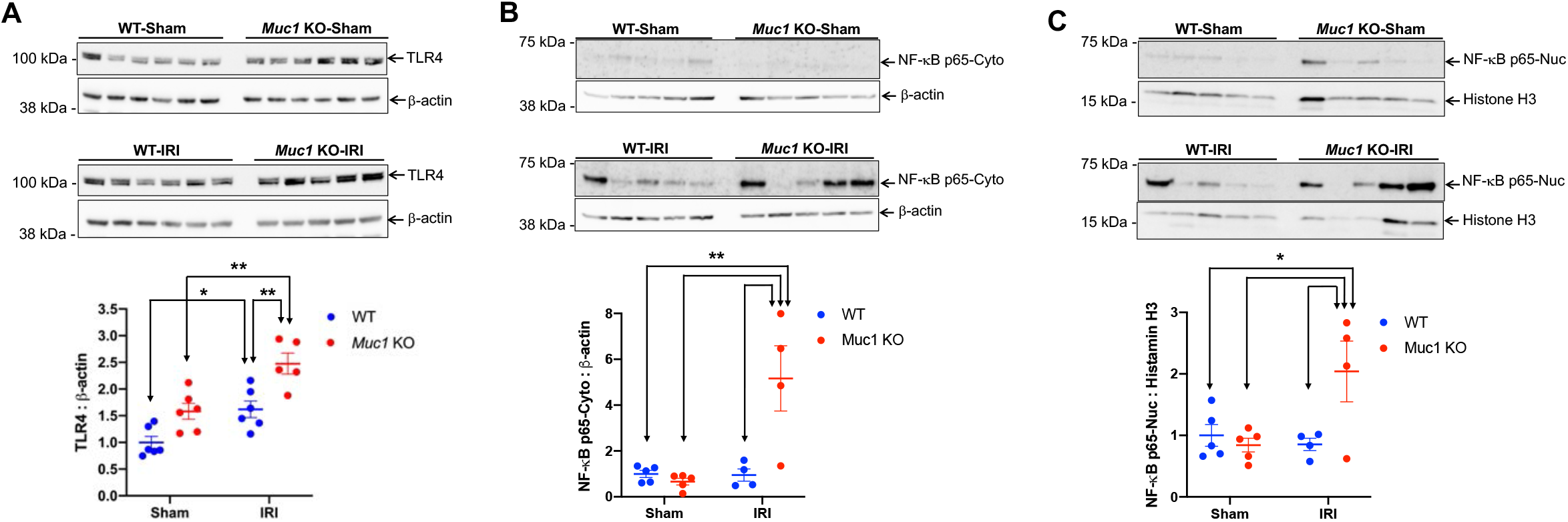
MUC1 stabilization of cell-associated KIM-1 correlates with reduced inflammation following IRI. Kidneys of *Muc1* KO mice (red) and wild-type (WT) littermates (blue) were subjected to 20 min ischemia and 48 h recovery (IRI) or sham surgery (Sham). (A) Kidneys were harvested at sacrifice and extracts of homogenates were subjected to immunoblotting for Toll-Like Receptor 4 (TLR4) and β-actin. (B-C) Aliquots of either nuclear extracts (Nuc) or cytoplasmic postnuclear supernatants (Cyto) of mouse kidneys were subjected to immunoblotting for NF-κB, and histone H3 or β-actin, respectively. TLR4 and NF-κB levels were normalized to β-actin or Histone H3 as indicated, and levels for WT-Sham were set to 1. Data were analyzed by two-way ANOVA (*: p<0.05; **: p<0.01; n=5-6).

**Figure 9.**
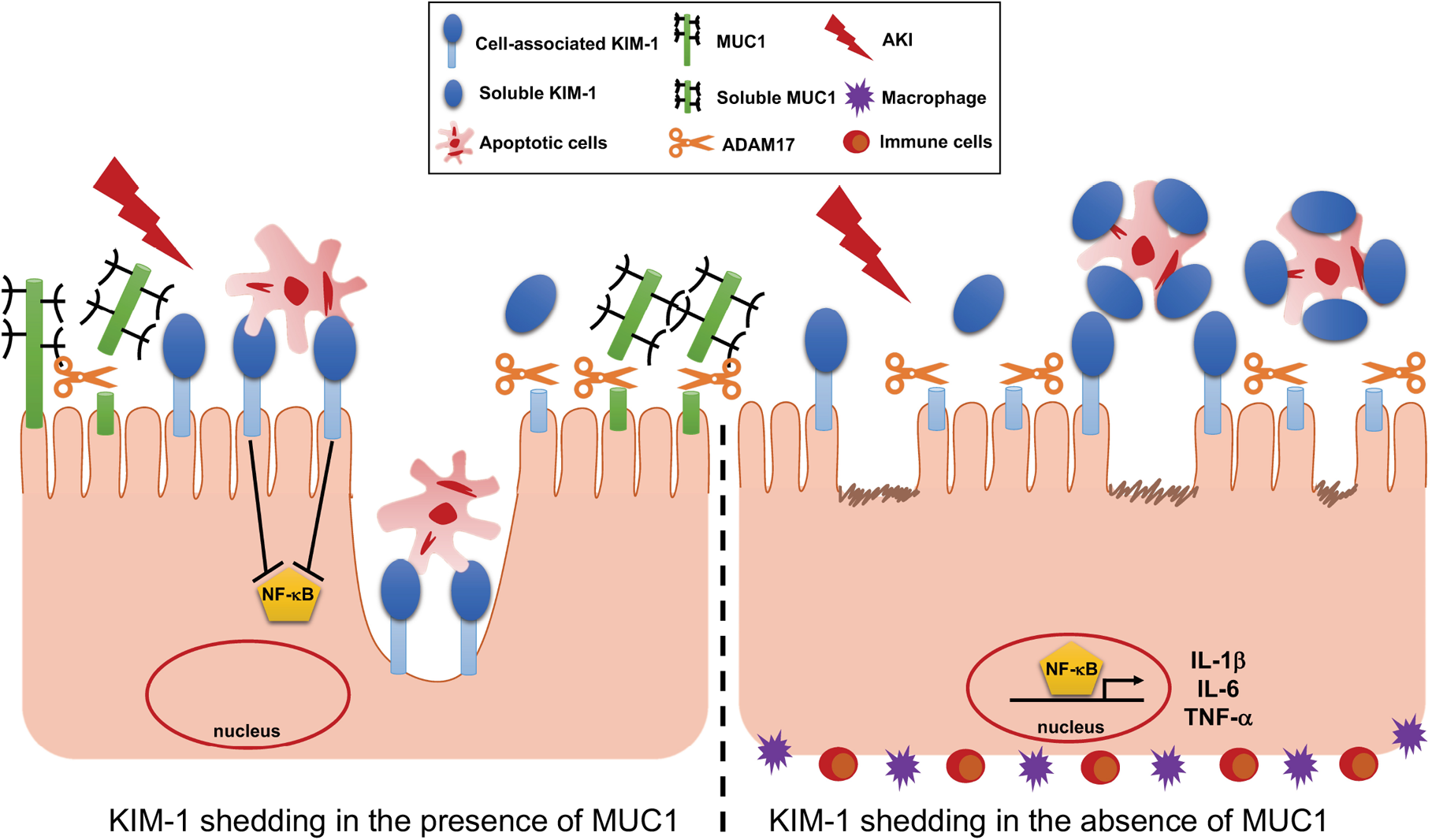
KIM-1 activity is preserved in the injured kidney by MUC1 inhibition of KIM-1 shedding. (LEFT) MUC1 induction in the PT following kidney injury reduces KIM-1 shedding by competing for ADAM17. (RIGHT) In the absence of MUC1, KIM-1 cleavage is accelerated and shed into the lumen where it acts as a decoy receptor, and subsequently inhibits efferocytosis. We hypothesize that stabilization of cell-associated KIM-1 by MUC1 induction during IRI could protect the kidney by maintaining KIM-1-mediated cell signaling and efferocytosis that reduces inflammation and promotes recovery.

## DISCUSSION

Using a mouse model of ischemia-reperfusion injury (IRI) where the renal pedicles were clamped for 19 min, we found that MUC1 predominately found in the TAL and later segments of the kidney tubule, was actually present and notably induced in the PT over 72 h recovery (17, 18). Kidney damage was worse, and recovery was blocked during IRI in *Muc1* KO mice when compared to congenic sham mice (based on histology and sCr), and we discovered that MUC1 stabilized both HIF-1α and β-catenin, their protective signaling pathways, and prevented metabolic stress including prolonged activation of AMPK (18, 19). In the present study, we used the hanging-weight protocol for renal IRI that has been shown to be a more clinically realistic model of IRI that provides highly reproducible kidney injury with virtually no tissue trauma or blood vessel congestion by blocking only the renal artery without venous occlusion (33, 34). After 20 min ischemia using this new protocol and 48 h recovery, we found that *Muc1* KO mice had more severe kidney injury than littermate WT mice, consistent with our previous work. We also found MUC1 induced in kidney extracts by immunoblotting, and MUC1 induced in all kidney tubules including the PT by confocal microscopy. While we were not surprised to find significantly higher levels of urinary KIM-1 in the *Muc1* KO mice when compared to WT mice due to the more severe kidney injury, we were surprised to find reduced levels of KIM-1 in the kidney extracts of the *Muc1* KO mice despite similar induction of mRNA levels in the Muc1 KO and WT kidneys. Our subsequent analysis was therefore focused on understanding how the absence of MUC1 could affect KIM-1 expression and consequently its function.

Given that KIM-1 is the most highly upregulated protein in the PT and because of its anti-inflammatory function following kidney injury (1, 2), there has been great effort to identify the upstream regulators of KIM-1. While results of two recent studies agree that phosphorylation of STAT3 is key to KIM-1 upregulation, two different kinases were implicated in rat (ChK1) and mouse (ERK1/2) studies (7, 8). However, we found no difference in STAT3 levels or activation between our sham and treated mice, and no difference between the *Muc1* KO and WT mice. While we could have missed any acute phase reactions by focusing on 48 h recovery (7, 8, 35), we also considered whether KIM-1 shedding was affected by the absence of MUC1.

We first asked whether MUC1 could increase KIM-1 shedding by regulating the expression of a relevant protease. ADAM17, also known as tumor necrosis factor (TNF-α) converting enzyme (TACE), is responsible for the ectodomain shedding of many cell surface proteins including KIM-1 and MUC1, and generation of many diverse active soluble ligands involved in development, regeneration, and inflammation (10, 14, 36, 37). In fact, Tang *et al*. found increased levels of active p38 MAPK and mature ADAM17 in a liver model of IRI (38), and Zhang *et al*. found that KIM-1 shedding was accelerated in a cell culture model by p38 MAPK activation (11). While we did find activation of p38 MAPK and increased ADAM17 in our mouse kidney model of IRI when compared to sham, we found no differences between *Muc1* KO and WT mice.

Alternatively, we asked whether MUC1 could compete with KIM-1 as a substrate for ADAM17. This possibility would be consistent with our observed increase in urinary levels observed in the absence of MUC1 during IRI in mice while kidney levels of KIM-1 were reduced. Using IF confocal microscopy of mouse and human kidneys, and an *in situ* proximity ligation assay in human kidneys, we first established that MUC1, KIM-1 and ADAM17 are co-localized at the apical plasma membrane of PTs. However, after ischemic injury, mouse KIM-1 is primarily associated with cellular debris in the tubule lumen in the absence of MUC1, suggesting that MUC1 blocks shedding of KIM-1. We also found that mouse KIM-1 shedding is stimulated in cultured MDCK cells (canine) by PMA, consistent with induction of ADAM17; and shedding is consistently reduced by co-expression with MUC1 and/or treatment of MDCK cells with a broad-spectrum metalloprotease inhibitor. Human MUC1 shedding is also reduced in primary cultures of transgenic mouse PT cells by an inhibitor specific for ADAM17 and ADAM10, but not by an inhibitor specific for ADAM10 alone. The consistency of our data implicating MUC1 inhibition of KIM-1 shedding is particularly interesting as human, murine and canine ADAM17 have greater than 90% amino acid sequence identity, while the transmembrane subunit of human and murine MUC1 exhibit only 65% identity that could reflect differences between species. Data from the Carson lab previously identified a specific site for human ADAM17 cleavage of human MUC1 at a position 58 residues from the transmembrane domain, but found no evidence of MUC1 cleavage by ADAMs 9, 10, 12 or 15 (14, 20). As recent studies have implicated ADAM10 in cleavage of human and mouse KIM-1 (12, 13), it becomes important to reassess the role of ADAM10 in mouse and human MUC1 shedding in future studies.

More importantly, we observed a difference in KIM-1 functional activity between *Muc1* KO mice and WT mice in kidneys during IRI. Cell-associated KIM-1 has an anti-inflammatory role as it mediates phagocytic clearance of apoptotic and necrotic cells (efferocytosis) following kidney injury, which in turn prevents inflammation and promotes kidney repair (1, 30). Furthermore, efferocytosis mediated by cell-associated KIM-1 also triggers KIM-1 signaling and thus downregulation of the pro-inflammatory NFκB signaling pathway (2). However, any accelerated shedding of KIM-1 competitively inhibits efferocytosis and its subsequent anti-inflammatory effects, as the excess soluble KIM-1 also acts as a decoy to cell-associated KIM-1 (10). Using confocal microscopy, we observed that KIM-1 was primarily apical (cell-associated) in the PT of WT mice where it mediates phagocytosis by interacting directly and partly surrounding the adjacent cellular debris following injury. In contrast, KIM-1 staining is confined mainly to the tubular lumen (cleaved form) together with cell debris even in the surviving PT of *Muc1* KO mouse kidneys after injury. Furthermore, using IF-staining, we also observed KIM-1 localized at the phagocytic cup and surrounding phagocytosed apoptotic cells within PTCs. More importantly, we noticed a significant difference in KIM-1 functional activity (efferocytosis) between *Muc1* KO and WT primary cultured PTCs. Finally, we demonstrated internalization of apoptotic cells within KIM-1 expressing cells but not adjacent KIM-1-negative primary cells.

Our findings are in agreement with a previous report in which authors confirmed the internalization of apoptotic cells by KIM-1 expressing cells (efferocytosis) when they demonstrated a direct contact between cell-associated KIM-1 and apoptotic cells using a confocal microscopy approach (1). Consistent with this distinct cellular distribution of KIM-1 after kidney injury, we found that kidneys of *Muc1* KO mice have more than a two-fold significant increase in active NF-κB levels (nuclear expression) than WT mice following IRI. Previous work showed a similar result in which greater inflammatory response and kidney dysfunction were reported in mice expressing KIM-1^Δmucin^ (KIM-1 protein missing the mucin domain required for apoptotic cells binding) compared to congenic shams following ischemic AKI (2). Altogether, these findings suggested that stabilizing cell-associated KIM-1 by MUC1 induction during IRI could protect the kidney by maintaining KIM-1-mediated efferocytosis and thus downregulating innate immunity and inflammation (Fig. 9).

## PERSPECTIVES AND SIGNIFICANCE

KIM-1 that is rapidly induced in the proximal tubule during renal IRI plays a key role in the recovery of the tubule epithelium by serving as a cell surface receptor for clearance of apoptotic cells (efferocytosis) and associated signaling that suppresses inflammation. While ADAM17 cleavage of KIM-1 yields a urinary biomarker, excessive cleavage provides a decoy receptor that aggravates efferocytosis and subsequent signaling. MUC1 is also induced during IRI and cleaved by ADAM17. Our data from studies in mice and cultured cells show that MUC1 competes with KIM-1 for cleavage by ADAM17. Consequently, MUC1 protects KIM-1 anti-inflammatory activity in the damaged kidney.

## ACKNOWLEDGMENTS

We thank Dr. Joseph Bonventre (Director, Renal Division, Brigham and Women’s Hospital, Harvard Medical School, Boston, MA) and Dr. Craig Brooks (Department of Cell and Developmental Biology, Vanderbilt School of Medical, Nashville, TN) for providing a cDNA for mouse KIM-1. We thank Dr. Olivera Finn (Department of Immunology, University of Pittsburgh School of Medicine, Pittsburgh, PA) for providing kidneys from transgenic MUC1 mice, and the Center for Organ Recovery and Education (CORE) for provision of human material for research (Pittsburgh, PA).

This work was supported by National Institutes of Health Grants K01DK109038, HL109002, DK091190, HL069846, DK070910, DK038470, the Kidney Imaging Core of the Pittsburgh Center for Kidney Research (P30-DK079307), and Dialysis Clinic, Inc.,

