## Supplemental Figures for "KIM-1-mediated anti-inflammatory activity is preserved by MUC1 induction in the proximal tubule during ischemia-reperfusion injury"

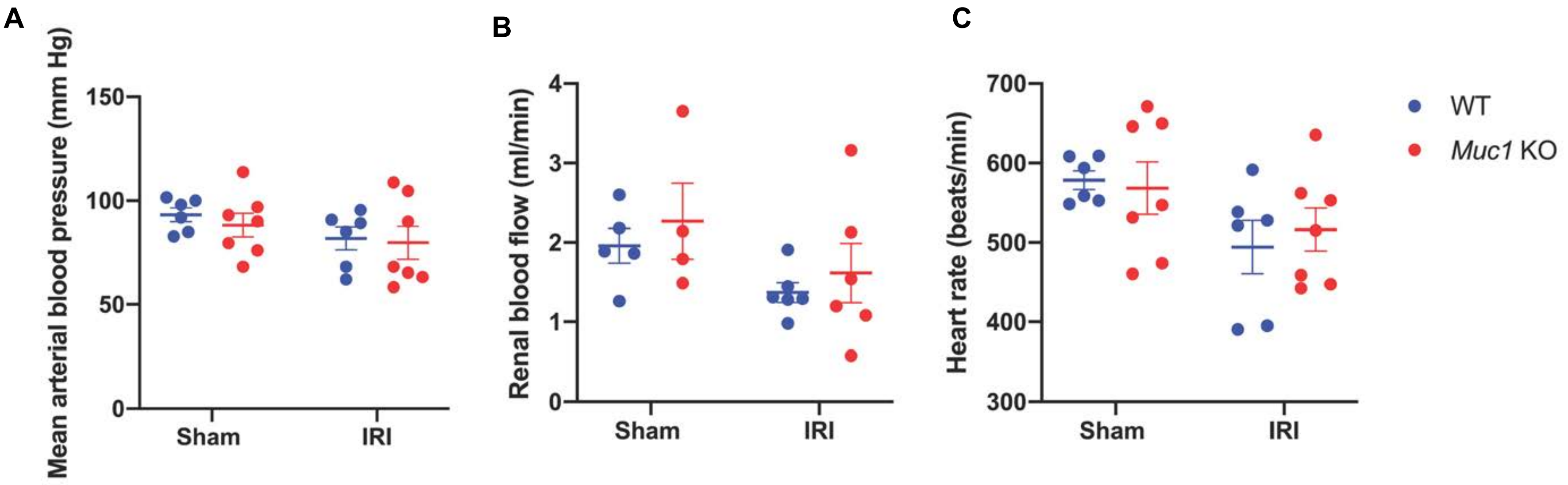


**Supplemental figure 1. No changes in kidney hemodynamics were observed during IRI.** Kidneys of *Muc1* KO mice (red) and wild-type (WT) littermates (blue) were subjected to 20 min ischemia and 48 h recovery using the kidney hanging-weight protocol (IRI) or sham surgery (Sham). Prior to sacrifice, mice were assessed for (A) mean arterial blood pressure, (B) renal blood flow, and (C) heart rate. Mean ± SE are shown. No significant differences were found by two-way ANOVA, (n=5-7).

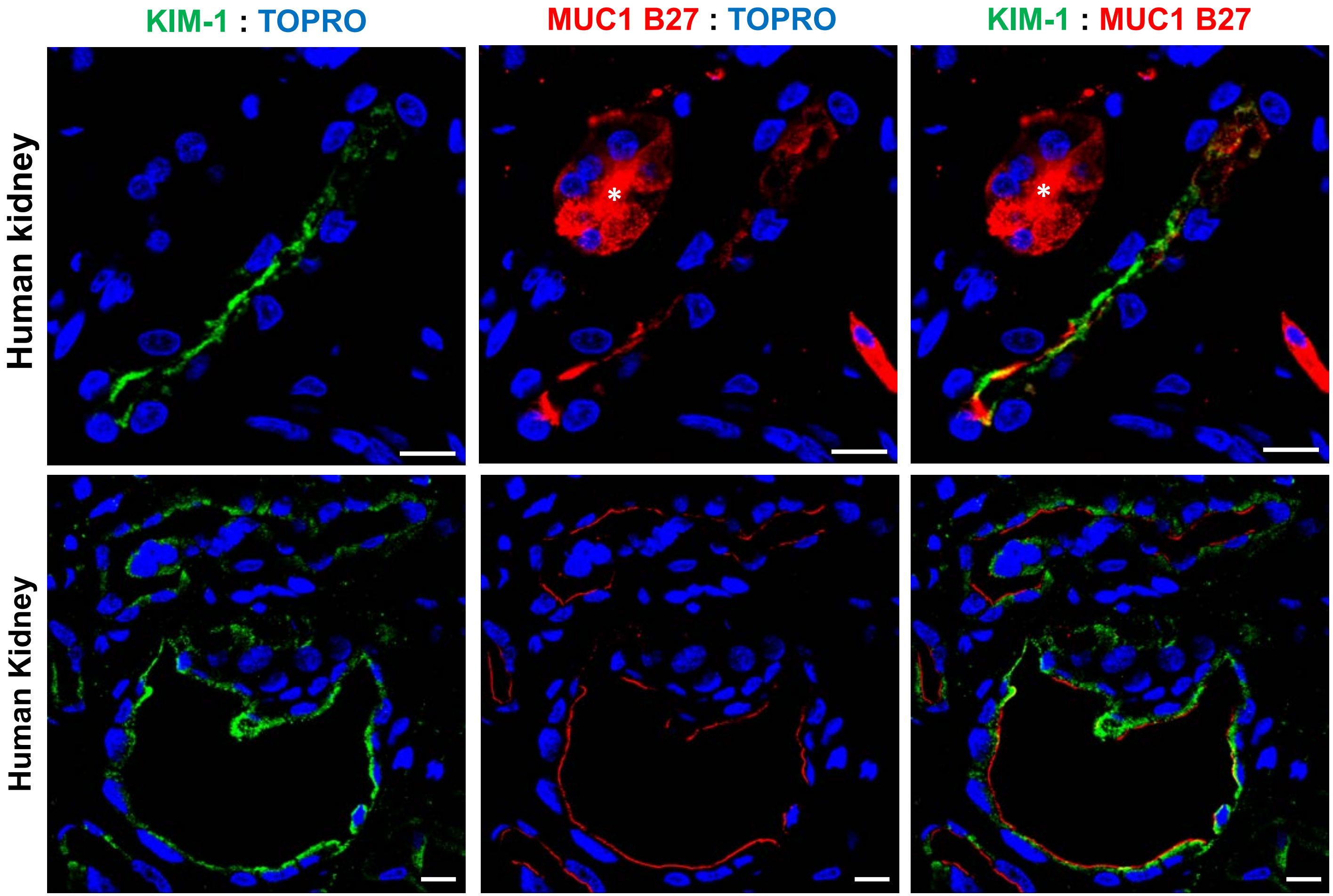


**Supplemental figure 2. KIM-1 and MUC1 are induced and co-expressed in human PT under ischemic conditions.** Human kidneys with ischemic damage were immunostained for KIM-1 (green), MUC1 (red) and nuclei (TOPRO, blue). Also, notice MUC1 expression in non-PT segments (labeled with an asterisk). Scale bar = 10 μm.

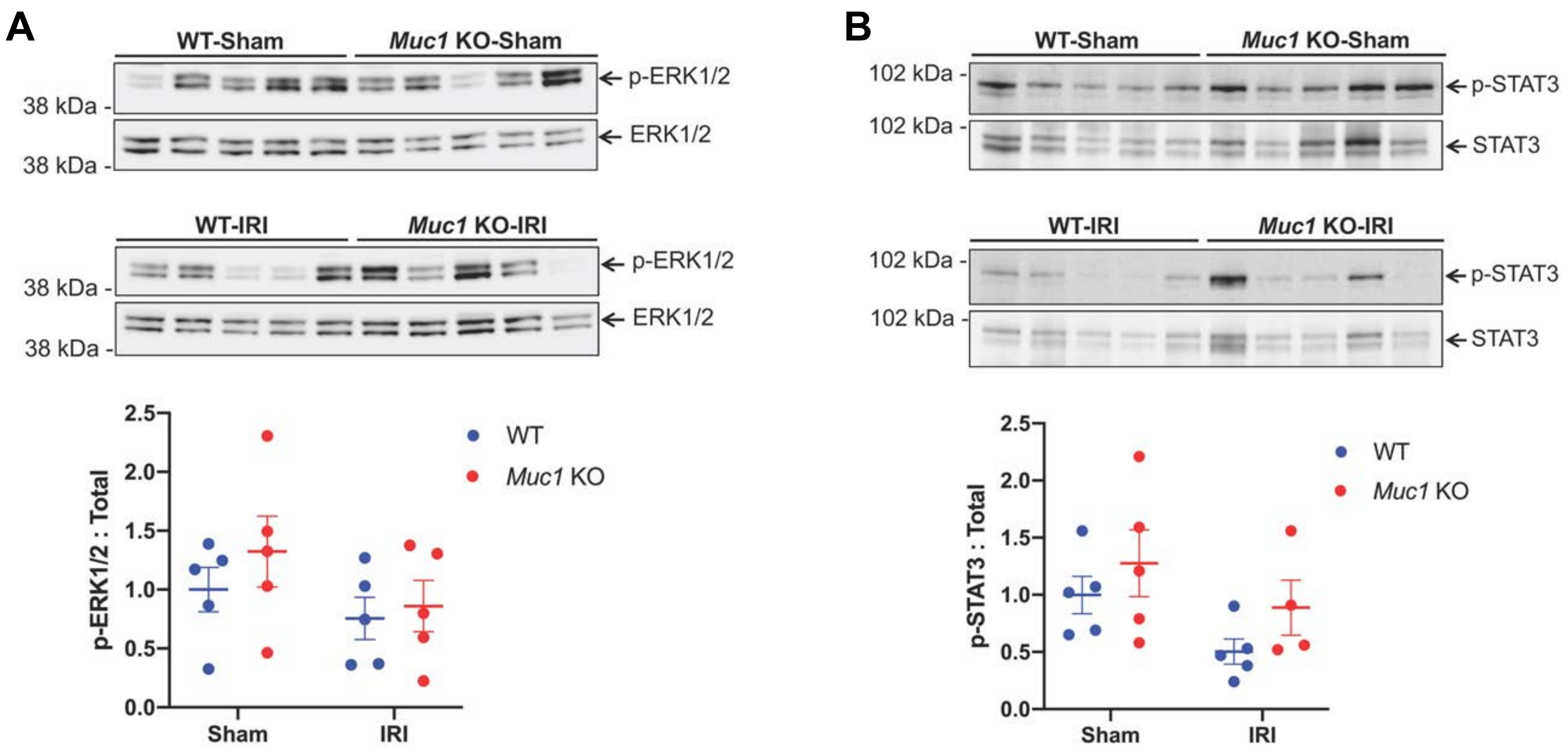


**Supplemental figure 3. No changes in the activation levels of upstream regulators of KIM-1 expression during IRI.** Kidneys of *Muc1* KO mice (red) and wild-type (WT) littermates (blue) were subjected to 20 min ischemia and 48 h recovery (IRI) or sham surgery (Sham). Kidneys were harvested at sacrifice and extracts of homogenates were subjected to immunoblotting for (A) p-ERK1/2 and ERK1/2, or (B) p-STAT3 and STAT3. The ratios of p-ERK1/2/total (A) and p-STAT3/total (B) are presented as means ± SE where the level for WT-Sham mice was set at 1. No significant differences were found by analyzing data with two-way ANOVA (n=5).

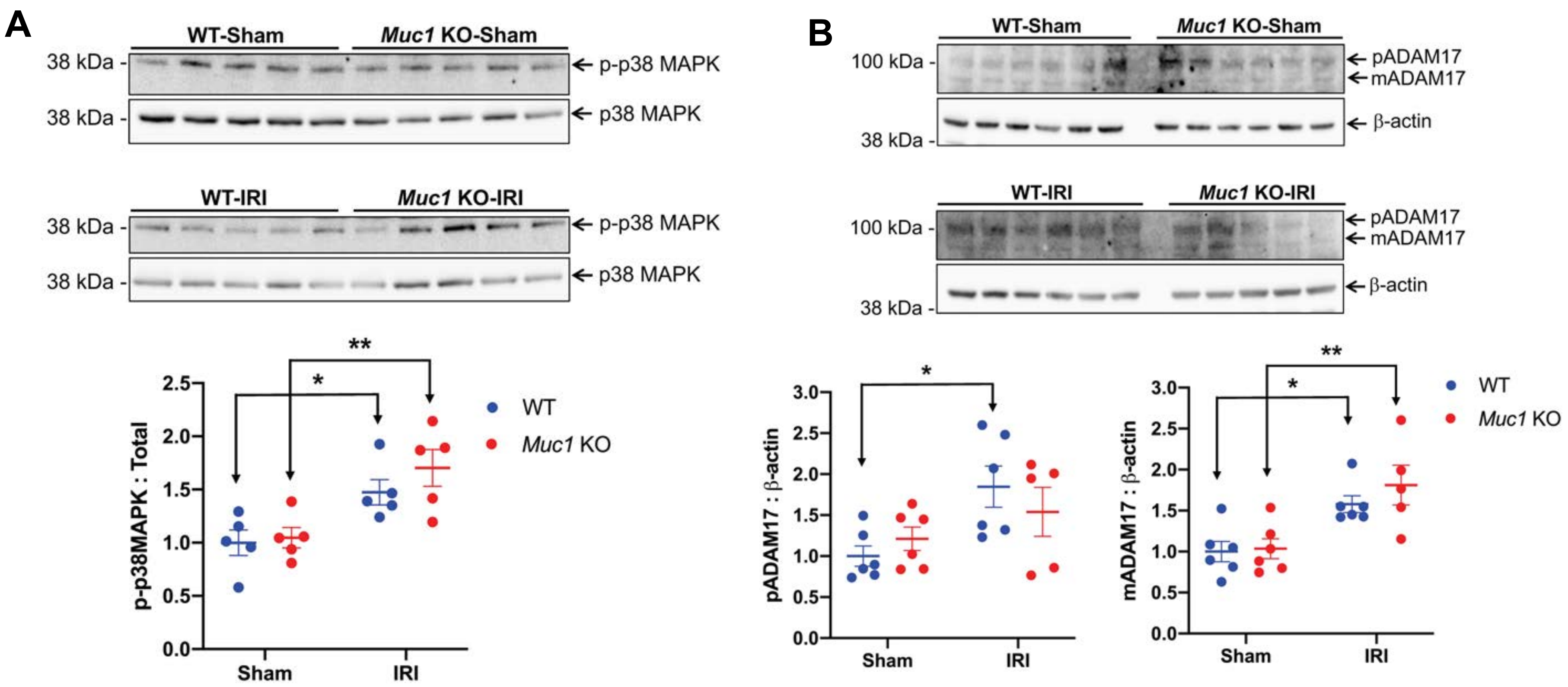


**Supplemental figure 4. Regulators of KIM-1 shedding are activated in the kidney during IRI.** Kidneys of *Muc1* KO mice (red) and wild-type (WT) littermates (blue) were subjected to 20 min ischemia and 48 h recovery (IRI) or sham surgery (Sham). Extracts of kidney homogenates were subjected to immunoblotting for (A) p-p38 MAPK and p38 MAPK or (B) ADAM17 and β-actin, as indicated. The p-p38 MAPK levels were normalized to total p38 MAPK, and ADAM17 levels were normalized to β-actin, while levels for WT-Sham are set to 1. Propetide ADAM17 (pADAM17) and mature cleaved ADAM17 (mADAM17) bands were analyzed separately. Data were analyzed by two-way ANOVA and significant differences are noted (*: $p<0.05$; **: $p<0.01$, n=5-6).

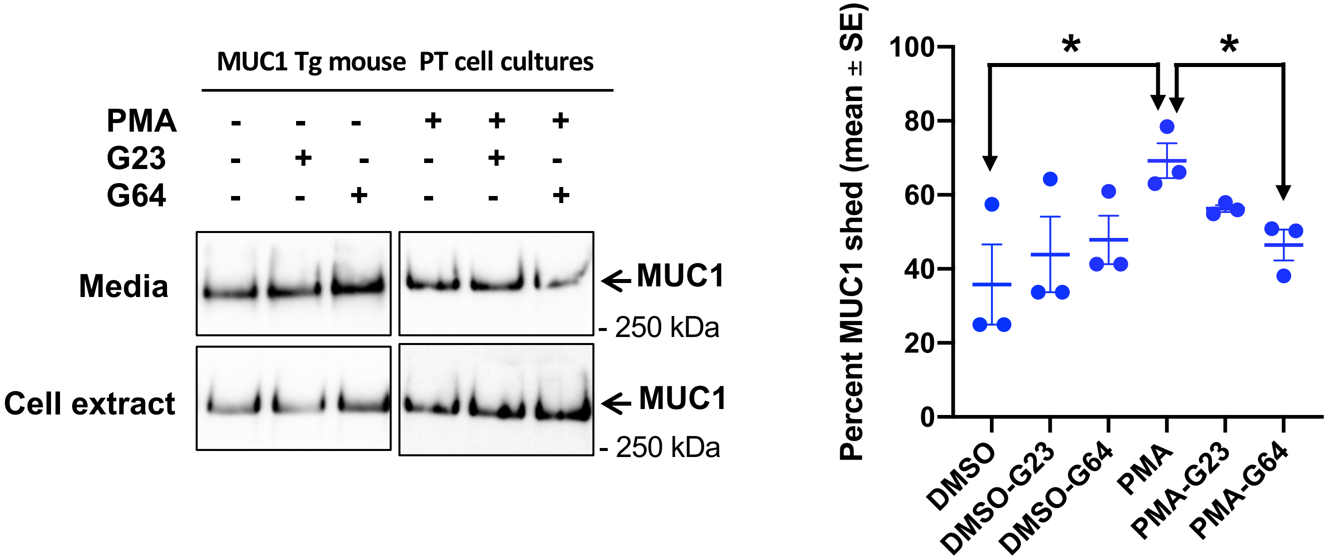


**Supplemental figure 5. MUC1 is shed from primary cultures of proximal tubule cells isolated from transgenic mice expressing human MUC1.** Primary cultures of proximal tubule cells from transgenic mice (n=3) were grown for one week on plastic, then treated with either PMA or vehicle (DMSO) in serum free media for 1 h in the presence or absence of protease inhibitors against both ADAM10 and ADAM17 (GW280264; **G64**) or specific for ADAM10 (GI254023; **G23**). Shedding of MUC1 was enhanced by treatment of cells with PMA that induces ADAM17, and was not blocked by specific ADAM10 inhibitor G23 but rather by the mixed ADAM17/ADAM10 inhibitor G64 indicating an important role of ADAM17 in MUC1 shedding. Data were analyzed by two-way ANOVA and significant differences are noted (*: $p<0.05$, n=3).

Supplementary Table 1. Primary antibodies used in this study.

| Target | Source | Host | Species reactivity (according to manufacturer) | Validation method |
|---|---|---|---|---|
| **MUC1 cytoplasmic tail (CT2)** | Gift from Sandra Gendler | Armenian hamster | Human, and Mouse | MUC1 CT KO mouse [18] |
| **MUC1 tandem repeat (TR)** | Fujirebio Diagnostics (B27.29) | Mouse | Human | MUC1 TR KO mouse [18-20] |
| **TIM-1/KIM-1** | R&D Systems (#AF1817) | Goat | Mouse | KIM-1 mucin domain deletion mice [21] |
| **TIM-1/KIM-1** | R&D Systems (#AF1750) | Goat | Human | Immunohistochemistry (company website) |
| **ADAM17** | Abcam (#ab2051) | Rabbit | Human, Mouse, and Rat | ADAM17 KD in HeLa cells (company website) |
| **STAT3** | Cell Signaling Technology (#9139) | Mouse | Human, Mouse, Rat, and Monkey | *STAT3 Knockdown & *Relative expression (company website) |
| **pS727-STAT3** | Cell Signaling Technology (#94994) | Rabbit | Human, Mouse, and Rat | Immunoprecipitation (company website) |
| **p38 MAPK** | Cell Signaling Technology (#8690) | Rabbit | Human, Mouse, Rat, Hamster, Monkey, Bovine, and Pig | *LLC-PK transfected with dominant negative p38 MAPK [22] *Immunohistochemistry (company website) |
| **p-P38 MAPK (Thr180/Tyr182)** | Cell Signaling Technology (#4511) | Rabbit | Human, Mouse, Rat, Monkey, Mink, Pig, and S. cerevisiae | LLC-PK transfected with dominant negative p38 MAPK [22] |
| **p44/42 MAPK (ERK1/2)** | Cell Signaling Technology (#9102) | Rabbit | Human, Mouse, Rat, Hamster, Monkey, Mink, Bovine, Pig, Zebrafish, and S. cerevisiae | ERK1/2 KO mouse [23] |
| **p-ERK1/2 (Thr202/Tyr204)** | Cell Signaling Technology (#9101) | Rabbit | Human, Mouse, Rat, Hamster, Monkey, Mink, Bovine, Pig, Zebrafish, D. melanogaster, and C. elegans | ERK1/2 KO mouse [23] |
| **TLR-4** | Cell Signaling Technology (#14358) | Rabbit | Mouse | 293T cells transfection with mock or TLR4 constructs (company website) |
| **NF-κB p65** | Cell Signaling Technology (#8242) | Rabbit | Human, Mouse, Rat, Hamster, Monkey, and Dog | Relative Expression (company website) |
| **OAT-1** | Alpha Diagnostic International (#OAT11-A) | Rabbit | Rat | Immunohistochemistry [24] |

Supplementary Table 2. Summary of patients with AKI enrolled in BioMaRK study.

| Patient # | Gender | Age | Serum creatinine (mg/dl) | Etiology | | | |
|---|---|---|---|---|---|---|---|
| | | | | Ischemic | Nephrotoxic | Sepsis | Multifactorial |
| 1 | female | 61 | 4.4 | Yes | Yes | Yes | Yes |
| 2 | male | 51 | 3.7 | Yes | No | Yes | Yes |
| 3 | female | 55 | 3.4 | Yes | Yes | Yes | Yes |
| 4 | male | 67 | 1.8 | Yes | No | Yes | Yes |
| 5 | female | 29 | 1.8 | No | No | No | Yes |
| 6 | male | 43 | 3.1 | Yes | No | Yes | Yes |
| 7 | female | 73 | 8 | Yes | Yes | Yes | Yes |
| 8 | male | 63 | 7.1 | Yes | No | No | Yes |
| 9 | female | 22 | 6 | No | Yes | No | Yes |
| 10 | female | 73 | 8.6 | Yes | Yes | Yes | Yes |

### Supplementary Table 3. Wedge biopsy reports of kidneys used in this study.

| Sample | Sclerotic Glomeruli | Interstitial Fibrosis | Arterial Nephrosclerosis | Arteriolonephrosclerosis | Glomerular Thrombi | Comments and Other Findings |
|---|---|---|---|---|---|---|
| HAK6R | 24 of 59 | Mild/Moderate | Mild/Moderate | Mild | None | Multifocal subcapsular and intraparenchymal interstitial chronic inflammation, Focal subcapsular scar |
| HAK30L | 2 of 105 | Mild | Mild | Mild | None | Foci of acute tubular necrosis about 5% |
| HAK31L | 61 of 117 | Mild/Moderate | Moderate | Mild/Moderate | None | Mesangial widening with relatively frequent Kimmelsteil-Wilson nodules, Patchy mild to moderate chronic interstitial inflammation |
| HAK63L | 4 of 53 | Mild | Mild | Mild | None | Occasional tubules with acute tubular necrosis, Multiple small foci of interstitial chronic inflammation |
